# A structural trade-off balances mechanical resilience and supramolecular adaptability in type IV pili

**DOI:** 10.64898/2026.07.02.736100

**Authors:** Chia-Ni Tsai, Laure Le Blanc, Fouad Mohamed Fouad Elgebely, Zainebe Al-Mayyah, Michael Nilges, Alexandre Persat, Yasaman Karami

## Abstract

Biological machines operate under mechanical load, requiring architectures that simultaneously resist force and remain functionally dynamic. Type IV pili (T4P) are bacterial filaments that experience large tensile forces during motor-driven retraction. Here, we combine cryo-electron microscopy, molecular dynamics simulations, optical tweezers, and functional analyses to define the structural basis of T4P mechanical adaptation. We determined a 2.8 Å cryo-EM structure of the *Pseudomonas aeruginosa* T4P and integrated it with comparative all-atom simulations across six bacterial strains to identify a conserved force-bearing electrostatic network. Simulations predicted that this network tunes filament elasticity under load, a finding validated by single-filament force spectroscopy. Rewiring these interactions experimentally produced hyper-rigid pili that assembled normally but exhibited impaired twitching motility. Together, our findings uncover a structural trade-off between force-resistant architecture and reversible supramolecular adaptability.

## Introduction

Bacteria rely on a wide range of surface-associated molecular machineries to sense, interact with, and adapt to their environments. Many of these systems evolved to perform highly specialized functions; for example, the bacterial flagellum primarily acts as a dedicated motility system that enables cells to navigate toward favorable environments and nutrients^1,2^. In contrast, type IV pili (T4P) are motorized surface filaments that support an exceptionally diverse repertoire of functions across bacterial species^3^. In pathogenic *Escherichia coli (E. coli*), T4P promote colonization of the intestinal mucosa^4^, whereas *Neisseria* spp. use them for host cell adherence and immune evasion^5,6^. *Myxococcus xanthus* relies on T4P for collective motility and predation^7^, and *Pseudomonas aeruginosa* utilizes T4P to drive twitching motility (**Fig. 1a**), mechanosensation, and biofilm development^8–15^. Thus, T4P can mediate both biochemical and mechanical functions, raising the question of how a single assembly with a highly conserved molecular architecture can simultaneously withstand large mechanical forces while supporting such diverse biological functions.

**Figure. 1:**
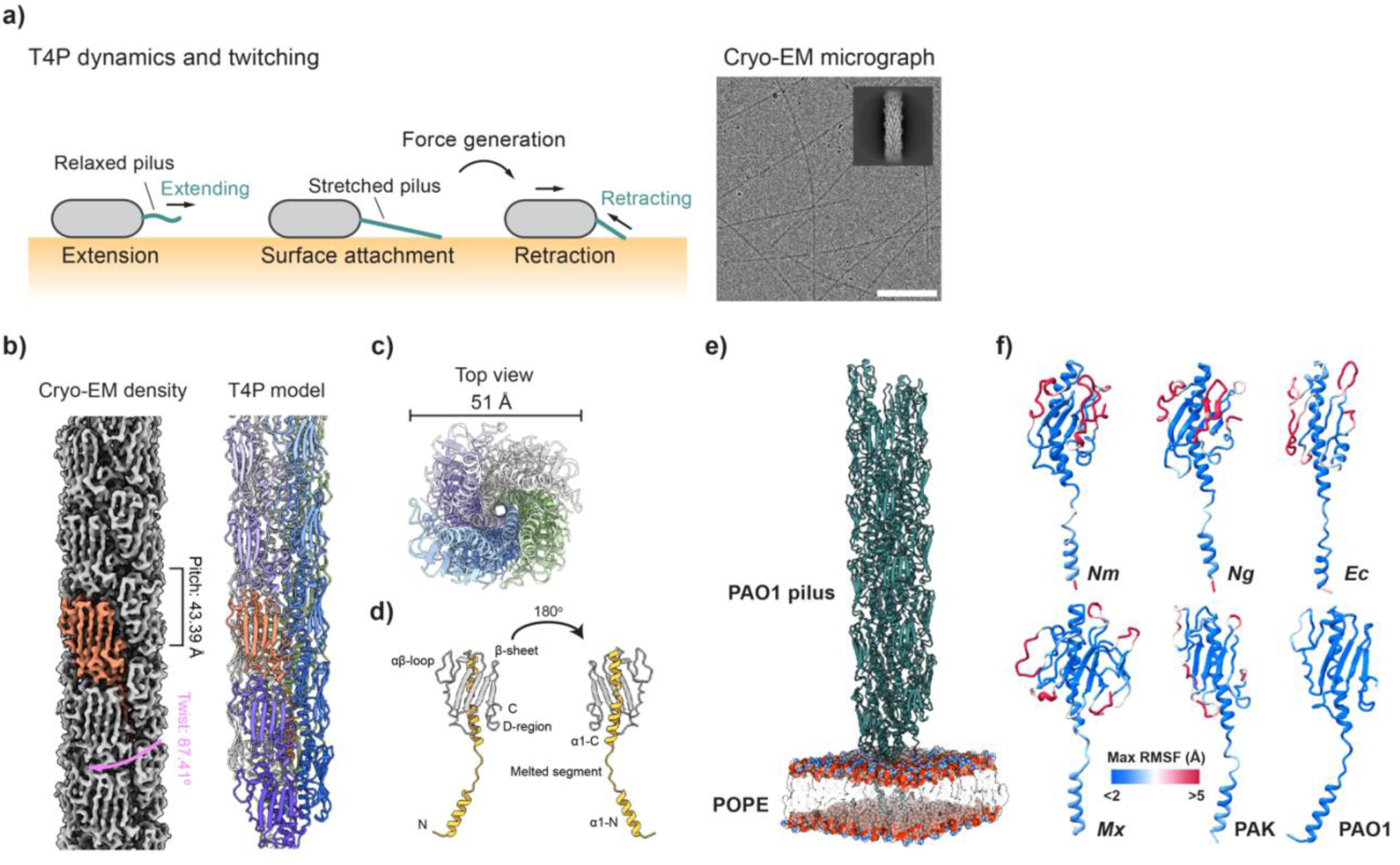
Equilibrium dynamics of T4P reflect functions across species. **a,** Schematic of twitching motility. The pilus (turquoise) extends from the cell envelope. Retraction of surface-attached pili generates force to drag the cell body toward the piliated pole. Cycles of extension and retraction drive twitching motility. Right: Representative cryo-EM micrograph of purified *P.* aeruginosa PAO1 T4P. Scale bar, 100 nm. The inset displays a representative 2D class average of vertically aligned filament particles, with visually distinguishable monomers. **b,** Left: Cryo-EM density map of the assembled PAO1 T4P reconstructed to 2.8 Å resolution. Right: Atomic model of the assembled PAO1 T4P. A single PilA subunit is highlighted in salmon in both panels. The filament exhibits a helical rise of 10.54 Å and a twist of 87.41°, corresponding to a helical pitch of 43.39 Å. **c,** Top view of the PAO1 T4P atomic model. The filament has an average diameter of ∼51 Å. The N-terminal α-helices pack tightly at the core, while the C-terminal globular domains face outward to form a hydrophilic surface. **d,** Structure of a single PAO1 PilA monomer. The N-terminal α-helix (α1) is highlighted in yellow and the C-terminal globular domain in white. The α1 helix is divided into α1-N and α1-C segments by an unstructured melted region. **e,** Initial structural model used for equilibrium molecular dynamics (MD) simulations. The PAO1 pilus is shown in cartoon representation (turquoise). The POPE membrane atoms are depicted as spheres and colored white (carbon), red (oxygen), and blue (nitrogen). **f,** Structural mapping of maximum root-mean-square fluctuation (RMSF) values onto individual major pilins from *N. meningitidis* (*Nm)*, *N. gonorrhoeae (Ng)*, *Ec*, *Mx*, PAK, and PAO1. The colormap indicates RMSF values from blue (rigid) to red (highly flexible). Values were calculated over the final 2.5 µs of n = 3 independent 3-µs MD simulations.

T4P are micrometers-long, nanometers-thin helical protein polymers of the major pilin subunits^16–18^. The ATPases PilB and PilT respectively drive pilin polymerization and depolymerization at the cell envelope to power T4P extension and retraction^19,20^. In *P. aeruginosa*, T4P extension, attachment, and retraction generate incremental displacements in a process called twitching motility^21–24^. In this process, attached T4P experience tensile forces of up to 100 pN generated by PilT, making it one of the strongest molecular motors in the living world^25^. Individual filaments must therefore maintain structural integrity under extreme tension while remaining capable of rapid extension and depolymerization, implying a finely balanced inter-subunit interaction network.

Despite their inherently dynamic functions, T4P have largely been investigated as static structures. As a result, the molecular mechanisms that preserve filament integrity under tension while enabling the structural adaptability required for repeated extension and retraction remain unclear. Biomechanical studies have nevertheless revealed key aspects of T4P mechanics. Atomic force microscopy (AFM) showed that *P. aeruginosa* T4P withstand forces up to 250 pN and adopt alternative conformations under tension^26^. Likewise, *N. gonorrhoeae* T4P undergo reversible conformational transitions under high pulling forces^27^. More recently, combining cryo-EM with biophysical measurements demonstrated that disrupting the inter-subunit interface of *M. xanthus* T4P changes filament persistence length, indicating that pilin packing governs filament mechanics^28^. Together, these studies suggest that T4P mechanics arise from a finely tuned structural trade-off between mechanical resilience and reversible structural rearrangements, yet the molecular interactions underlying this balance remain unknown.

Establishing causal links between atomic-level interactions and emergent filament mechanics remains challenging. Equilibrium molecular dynamics (MD) predicts how local structural features shape force propagation and mechanical behavior, and we previously used it to identify hydrogen bonds and salt bridges governing the stability and flexibility of the *E. coli* T4P^29^. However, because equilibrium MD does not capture externally applied forces, it cannot explain how T4P respond to mechanical loading. Steered MD (SMD) addresses this limitation by simulating protein deformation under force. Combined with AFM, SMD has revealed the molecular basis of the remarkable resilience of type I pili^30,31^. Previous atomistic and coarse-grained SMD studies likewise investigated T4P mechanics but relied on structural models containing a continuous N-terminal α-helix, limiting insight into the force-bearing interactions of native pili^32,33^. The recent availability of near-atomic cryo-EM structures now enables high-resolution simulations of how T4P reconcile two competing mechanical requirements: maintaining filament integrity under tension while preserving the reversible structural rearrangements required for pilin extraction during retraction. Combined with single-molecule force measurements, this approach provides a powerful framework for linking atomistic interactions to the emergent mechanics of T4P.

Here, we combine molecular dynamics simulations, single-filament biophysical measurements, and mutational analysis to investigate the mechanical behavior of T4P across six Gram-negative bacteria: *Neisseria meningitidis (Nm)*, *N. gonorrhoeae (Ng)*, *Escherichia coli (Ec)*, *Myxococcus xanthus (Mx)*, and *Pseudomonas aeruginosa* (PAK and PAO1). By linking atomistic interactions to filament-scale mechanics, we uncover a conserved force-bearing interaction network and show that T4P function depends on a structural trade-off between mechanical resilience and reversible supramolecular adaptability.

## Results

### Comparative MD reveals distinct intrinsic flexibilities of T4P pilins

To investigate the intrinsic stability of T4P, we performed comparative molecular dynamics (MD) simulations of six T4P systems from different bacterial species (three replicates of 3-μs simulations per system, *Ec* simulations from reference^29^). Because precise structural coordinates and side-chain orientations are prerequisites for accurate atomistic simulations, we determined a higher resolution structure of the intact PAO1 T4P using cryo-EM. We employed a two-step refinement strategy to achieve a high-resolution reconstruction. First, cryo-EM micrographs (**Fig. 1a, right**) of purified pili underwent filament tracing and iterative helical real-space refinement, yielding an initial map at 3.1 Å resolution. Because strict global symmetry can blur less ordered regions such as loops and linkers, we subsequently applied non-uniform refinement to the helical map. This adaptive regularization down-weighted poorly ordered regions while reinforcing the well-aligned core, preserving high-resolution features and improving the final overall resolution to 2.8 Å (**Fig. 1b**). This resolution permitted the unambiguous assignment of side-chain conformations required for downstream computational analyses. In this optimized map, the major pilin PilA monomer (salmon) and its surface-exposed β-sheets are clearly resolved. From this map, we constructed an atomic model containing 22 subunits, revealing a filament diameter of ∼51 Å (**Fig. 1c**). We note that recently published structures of the PAO1 T4P at 3.9 and 3.2 Å resolutions align closely with our data^18,34^. In addition, we used previously published T4P structures from *Nm*, *Ng*, *Ec*, *Mx*, and PAK^17,28,35,36^ (**Extended Data Table 1**).

For all six strains, pilin monomers extracted from the T4P structures exhibit a conserved architecture consisting of an N-terminal α-helix (α1), an αβ-loop connecting α1 to a globular domain formed by an antiparallel β-sheet, and a C-terminal D-region (**Fig. 1d**). Despite a conserved global helical organization, the major pilins from *Nm*, *Ng*, *Ec*, *Mx*, PAK, and PAO1 differ in several topological features, including variations in C-terminal β-sheet extensions, short accessory helices, and disulfide-bond patterns. To determine how these species-specific structural variations influence global filament stability, we evaluated the baseline fluctuations of individual pilin subunits across all six models from their equilibrium MD simulations (**Fig. 1e**). To quantify structural dynamics, we calculated the root-mean-square fluctuation (RMSF) of the backbone atoms (*Cα, C, N,* and *O*) for each residue relative to the time-averaged conformation over the last 2.5 µs of the 3-µs MD simulations. Mapping the maximum RMSF values onto individual pilin subunits shows that structural fluctuations are concentrated in loop regions (**Fig. 1f**). *Nm* and *Ng* exhibited the highest mean RMSF values (1.79 ± 1.41 Å and 1.65 ± 1.15 Å, respectively). *Ec* and *Mx* demonstrated intermediate flexibility (1.49 ± 0.73 Å and 1.48 ± 1.01 Å, respectively), whereas PAK and PAO1 proved the most rigid, displaying the lowest mean RMSF values (1.18 ± 0.45 Å and 0.77 ± 0.32 Å, respectively).

We hypothesized that inter-subunit interactions contribute to T4P conformational stability during equilibrium MD. To identify these interactions, we quantified hydrogen bonds and salt bridges between neighboring pilins across all six models (**Extended Data Fig. 1**). Most systems maintained multiple inter-subunit interactions involving surface-exposed loop residues, indicating that these flexible regions also contribute to filament stabilization. An exception was PAK, which formed only a single loop-mediated hydrogen bond (K110–D124). Across all models, we identified a conserved electrostatic interaction linking adjacent pilins (R30–E49 in *Nm*, *Ng*, PAK, and PAO1; K30–E53 in *Ec*; and R30–E53 in *Mx*), in which a positively charged lysine or arginine simultaneously forms a hydrogen bond and salt bridge with a glutamate residue. This conserved interaction likely stabilizes the hydrophobic core of the assembled filament. We also observed the previously described T2–E5 hydrogen bond between adjacent subunits in *Nm*, *Ng*, *Ec*, and PAK^17,36–39^, supporting its proposed role in pilus assembly. In contrast, this interaction was absent from both the initial structures and equilibrium MD trajectories of *Mx* and PAO1, indicating that these species rely on distinct inter-subunit interaction networks.

### SMD reveals divergent force responses of assembled T4P

We next investigated how inter-subunit interactions preserve filament integrity under load by conducting constant-velocity steered molecular dynamics (SMD) simulations on T4P (five replicates of 200 ns simulations per system). A stretching force was applied to the center of mass of the four top subunits (rate of 1 Å/ns, spring constant 1000 kJ mol⁻¹ nm⁻²), while the positions of the first 30 residues of the four bottom-most subunits were fixed (**Fig. 2a**). To quantify force propagation over time, we analyzed the bulk section of the filament, excluding the top and bottom four subunits. Across all six homologs, the pulling force increased rapidly during the initial tens of nanoseconds before gradually decaying (**Fig. 2b**). Among the six T4P models, PAO1 T4P exhibited the highest maximum restoring force and maintained the largest residual force at the end of the simulations, whereas the other systems relaxed to near-zero values.

**Fig. 2:**
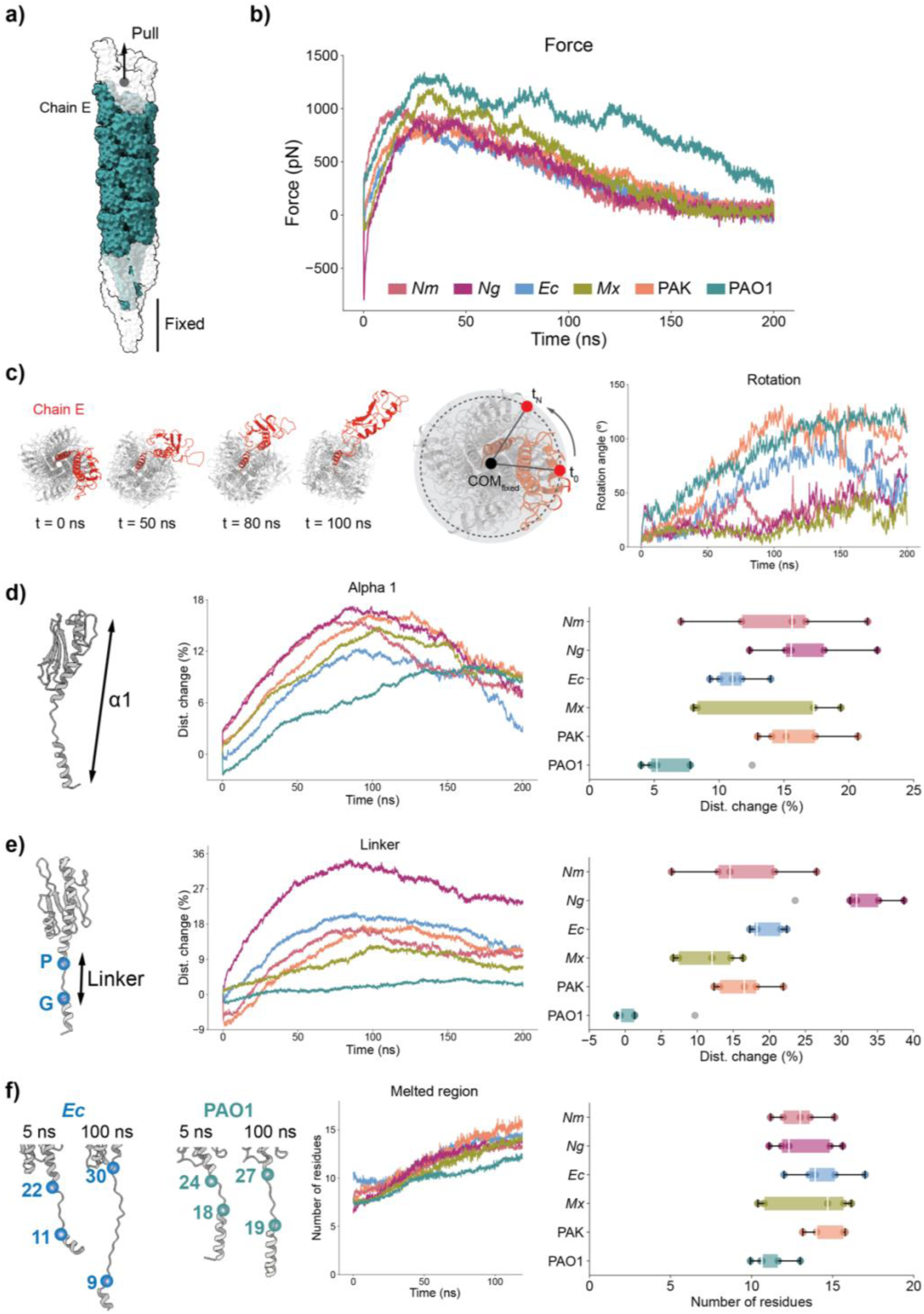
Steered molecular dynamics (SMD) simulations and quantification of the T4P force response. **a,** Structural model of the *P. aeruginosa* PAO1 T4P filament used for constant-velocity SMD simulations. The “bulk” subunits are highlighted in turquoise, while the four top-most and four bottom-most subunits are shown in white. Pulling force was applied to the center of mass of the four top subunits (gray circle). The N-terminal 30 residues of the four bottom-most subunits were position-restrained during the simulations. POPE model membrane is not shown. **b,** Temporal evolution of the pulling force applied to the T4P filaments. The species color code applies to all subsequent panels. Curves represent the mean; *n* = 5 independent 200-ns replicates per species. **c,** Force-induced rotational dynamics of T4P. Left: Top-view snapshots of PAO1 T4P at 0, 50, 80, and 100 ns. Structures are aligned by their fixed bottom subunits. Chain E is highlighted in red to illustrate rotational motion. Middle: Schematic defining the rotation angle, measured between the center of mass of chain E at *t =* 0 and *t = n* (red dots), using the center of mass of the fixed bottom residues (COM_fixed_) as the pivot. Right: Temporal evolution of the rotation angle. Values represent the mean rotation of chain E; *n* = 5 independent replicates. **d,** Mechanical extension of the pilin core α1 helix. Left: Initial structure of the PAO1 major pilin monomer. The arrow indicates the measured distance of the α1 helix. Middle: Time evolution of the normalized change in α1 helix length (mechanical tensile strain). Right: Distribution of α1 mechanical strain values at 100 ns across the bulk subunits; *n* = 5 replicates. **e,** Mechanical extension of the pilin linker region. Left: Initial structure of the *E. coli* (*Ec*) major pilin monomer. Conserved glycine and proline residues flanking the linker are shown as blue spheres; the arrow indicates measured linker distance. Middle: Time evolution of the normalized change in linker length. Right: Distribution of linker mechanical strain values at 100 ns across bulk subunits; *n* = 5 replicates. **f,** Force-induced unfolding of the α1 helix melted regions. Left: Initial structures of *Ec* and PAO1 major pilin monomers. Flanking residues of the melted regions are depicted as blue and turquoise spheres, respectively. Middle: Time evolution of the number of unstructured (non-α-helical) residues in the melted segments, determined by DSSP analysis. Only the first 120 ns of the 200-ns simulation are shown; the full 200-ns curve is provided in **Extended Data Fig. 2a**. Right: Distribution of the number of melted residues at 100 ns across bulk subunits; *n* = 5 replicates. For all box plots in d–f, the center line indicates the median, box limits represent the upper and lower quartiles, and whiskers indicate 1.5× the interquartile range (IQR). Outliers are shown in gray.

The SMD simulations revealed pronounced helical untwisting during filament stretching (**Fig. 2c, left panels**). We monitored the rotation of chain E, the first subunit adjacent to the region where force was applied. To quantify this rotational behavior, we used the center of mass of the restrained bottom subunits (COM_fixed_) as a pivot (**Fig. 2c, middle**). We tracked the angular deviation of chain E from its starting position over time and averaged this measurement across the five replicates. The magnitude of stretch-induced untwisting differed across species: PAK and PAO1 exhibited the largest angular changes, *Ec* showed an intermediate rotation, and *Nm*, *Ng*, and *Mx* rotated the least. Combined with our equilibrium MD data, these results demonstrate that the PAO1 T4P is the most mechanically stable filament prior to force application and maintains the highest structural rigidity under sustained tension.

We next examined the mechanical response of individual pilin subunits during stretching. Because the N-terminal α1 helix and internal linker region form the structural core of T4P, we quantified their force-induced extension throughout the SMD simulations (**Fig. 2d, e**). The linker region is defined as the segment between a conserved glycine and proline, whose initial length and specific residue boundaries (e.g., G14–P22 or G11–P22) differ across systems. Most systems exhibited a biphasic response characterized by initial elongation followed by partial relaxation. In contrast, PAO1 pilin consistently displayed the lowest deformation. At 100 ns, the length of the α1 helix of *Nm, Ng, Ec, Mx,* and PAK extended by ∼11–15%, whereas PAO1 extended by only 5% (**Fig. 2d**). Similarly, while linker regions in the other species elongated by 13–30% of their resting lengths, the PAO1 linker remained essentially unchanged throughout the simulation (**Fig. 2e**). Together, these observations indicate that the PAO1 pilus core is markedly less extensible under load than those of the other species.

Pilins contain a partially melted hinge region within the α1 helix that is thought to provide the flexibility required for filament assembly and deformation. Here, the melted region is defined as the unstructured segment lacking α-helical content. To quantify force-induced structural rearrangements, we monitored the α1 secondary structure throughout the SMD simulations using DSSP^40^ (**Fig. 2f and Extended Data Fig. 2a**). All systems exhibited dynamic unfolding and refolding under tension, consistent with ongoing local remodeling of the hinge region. However, PAO1 pilin consistently displayed the fewest unstructured residues and the smallest increase in α1 melting during stretching (**Extended Data Fig. 2b**), indicating enhanced resistance to force-induced deformation.

Together, these simulations show that PAO1 T4P possesses the most rigid pilin architecture among the studied systems, while retaining structural adaptability through enhanced filament untwisting. Analysis of the hydrogen-bond and salt-bridge networks during SMD simulations revealed that, although most interactions were disrupted by stretching, the T2–E5 hydrogen bond and a conserved inter-subunit electrostatic interaction (R30/K30–E49/E53) remained stable under tension across most of the species (**Fig. 3a and Extended Data Fig. 3**). These interactions define a conserved force-bearing network that maintains filament integrity during deformation. We therefore experimentally tested their contribution to T4P mechanics and biological function.

**Fig. 3:**
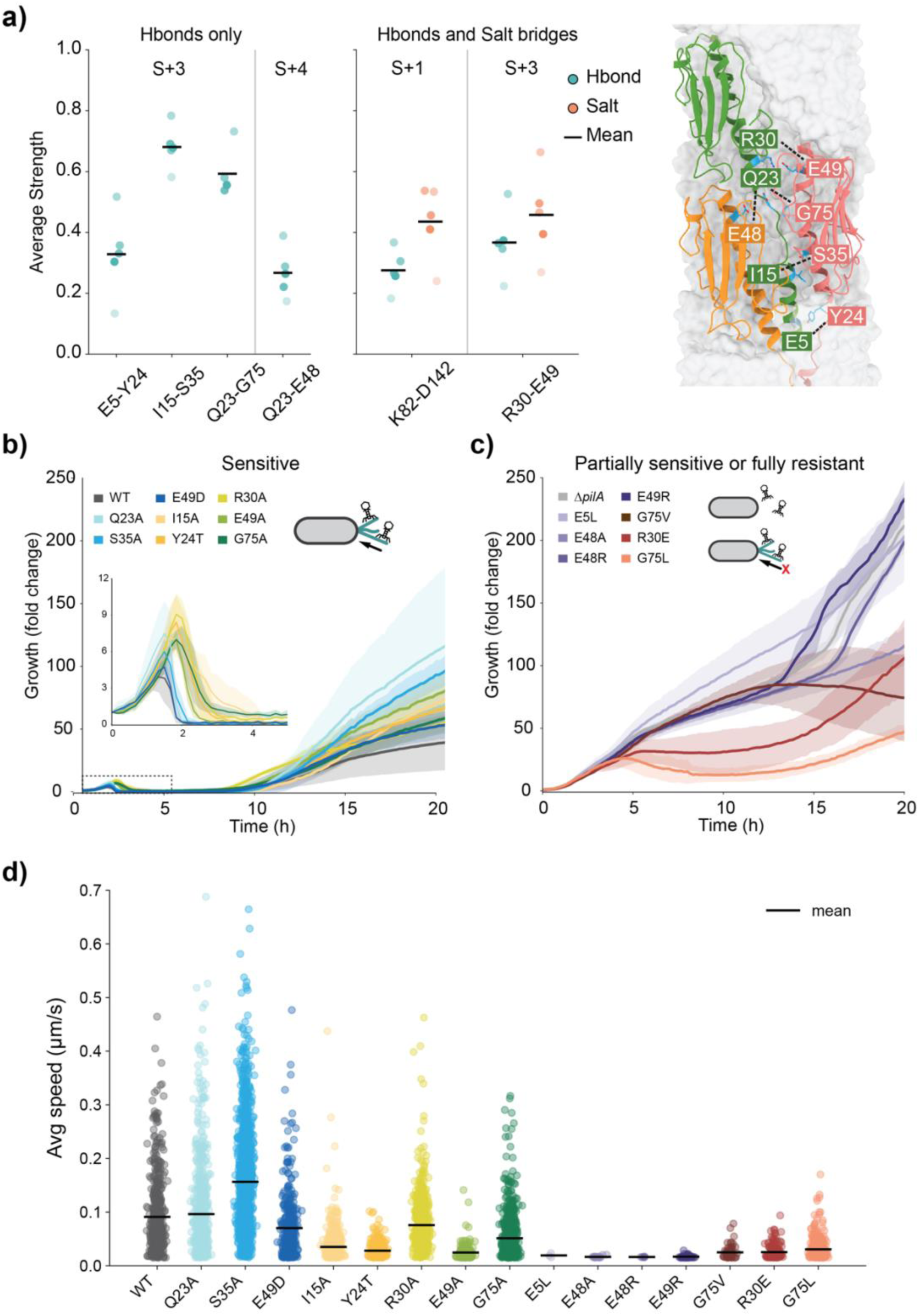
Force-resistant inter-subunit interactions govern T4P assembly function in *P. aeruginosa* PAO1. **a,** Inter-subunit interaction networks of PAO1 T4P. Left: Interaction networks characterized exclusively by hydrogen bonds (Hbond, turquoise). Middle: Interaction networks involving both hydrogen bonds and salt bridges (salmon). Each circle represents the mean contact persistence from a single steered molecular dynamics (SMD) replicate; the horizontal black line indicates the overall mean; *n* = 5 independent replicates. Right: Ribbon representation of three adjacent pilin subunits (S, yellow; S_+1_, pink; S_+4_, green). The structural pairs targeted for mutagenesis, corresponding to the left and middle panels, are connected by black lines. **b,** Phage infection growth curves for WT PAO1 and fully phage-sensitive PilA mutants. Bacterial density (fold-change relative to initial concentration) was monitored over 20 h following infection with the T4P-dependent lytic phage LUZ19. WT PAO1 and fully sensitive mutants exhibiting nearly identical growth and lysis kinetics are shown in blue shades. Mutants displaying slightly delayed lysis and marginally higher persistent cell densities are shown in yellow and green. The schematic illustrates successful phage binding to, and retraction with, a functional T4P. **c,** Phage infection growth curves for fully resistant and partially sensitive PilA mutants. Fully resistant strains, exhibiting continuous exponential growth without lysis, are shown in gray and purple shades. A severely impaired mutant (G75V) is shown in brown. Partially sensitive mutants are shown in red (R30E) and orange (G75L). The schematic illustrates the failure of phage infection due to absent or non-functional T4P. For simplicity in the plot legends in b and c, WT refers to the *ΔfliC* parental strain, and all *ΔpilA* and *pilA* mutants were generated in this *ΔfliC* background. In b and c, solid lines represent the mean; shaded areas represent s.d.; *n* = 3 biological replicates. **d,** Single cell twitching motility assay and average speed analysis. Non-motile cells were excluded from the velocity plots. Each circle represents the average twitching speed of an individual tracked cell. Black horizontal lines indicate the mean of the average speeds for each strain. The color palette corresponds to the functional groups defined in b and c. The percentage of actively twitching cells for each strain is as follows: WT (89.8%, 543/605), Q23A (90.9%, 683/751), S35A (98.8%, 1360/1376), E49D (96.8%, 387/400), I15A (82.7%, 966/1168), Y24T (37.2%, 312/839), R30A (96.1%, 708/737), E49A (22.7%, 222/979), G75A (89.7%, 833/929), E5L (0.4%, 3/731), E48A (3.5%, 18/514), E48R (0.7%, 5/706), E49R (10.5%, 52/495), G75V (7.2%, 62/866), R30E (26.9%, 187/695), and G75L (43.2%, 433/1002). For each strain, values in parentheses denote the percentage of motile cells, followed by the raw counts representing (motile tracks / total tracks)

### Conserved inter-subunit interactions stabilize T4P under tensile load

To investigate the functional contribution of inter-subunit interactions, we generated targeted *pilA* point mutants in PAO1. Guided by the equilibrium MD and SMD simulations, we selected five predicted interaction pairs for mutagenesis: E5–Y24, I15–S35, Q23–G75, Q23–E48, and R30–E49 (**Fig. 3a**). We assessed T4P function using the lytic phage LUZ19, which requires functional pili for infection^41^. As expected, wild-type (WT) cells underwent complete lysis following an initial growth phase, whereas the Δ*pilA* strain remained fully resistant, establishing a clear distinction between functional and non-functional pili (**Fig. 3b, c**). Mutating the highly conserved E5 residue produced a fully resistant phenotype comparable to Δ*pilA*, indicating complete loss of T4P function. In contrast, mutation of its predicted interaction partner Y24 had only a minor effect on phage susceptibility. A similar asymmetry was observed for the Q23–E48 pair: whereas E48 was essential for T4P function, mutation of Q23 had little effect on infection kinetics. These results suggest that the critical roles of E5 and E48 arise from additional structural interactions beyond their predicted binding partners. When these primary partners (Y24 or Q23) are mutated, E5 and E48 likely form novel compensatory interactions to maintain functional T4P assembly.

To assess the functional importance of the conserved R30–E49 interaction, we systematically perturbed its electrostatic properties. Introducing a charge-reversal mutation at position 49 (E49R) completely abolished phage infection, indicating a loss of T4P function. In contrast, a charge-conserving substitution (E49D) retained full infectivity. Mutation of the complementary residue (R30E) produced an intermediate phenotype: infected cultures reached a higher initial density than WT and underwent delayed, partial lysis, consistent with the formation of partially functional pili. Together, these results demonstrate that precise electrostatic complementarity at the conserved R30–E49 interface is required for optimal T4P function *in vivo*.

We also observed a striking side-chain-dependent modulation of T4P function at position G75. Substitution with alanine (G75A) had no detectable effect, whereas the slightly larger valine substitution (G75V) almost completely abolished T4P function, producing only minimal lysis at late stages of phage infection. Unexpectedly, the bulkier leucine substitution (G75L) partially restored function, resulting in delayed lysis following initial cell growth. This non-monotonic response indicates that T4P function is highly sensitive to the local packing environment surrounding G75 and suggests that the leucine side chain can establish compensatory interactions that are inaccessible to valine. Control experiments confirmed that the altered infection phenotypes of the R30 and G75 mutants were not attributable to general growth defects (**Extended Data Fig. 4**).

We next investigated how these inter-subunit interactions influence T4P-dependent twitching motility, a process driven by pilus retraction under piconewton-scale tension. Single-cell tracking showed that ∼90% of WT PAO1 cells were motile (**Fig. 3d**). Consistent with the phage infection assays, mutants that completely lost phage sensitivity (E5L, E48A, E48R, and E49R) were essentially non-motile. In contrast, phage-sensitive mutants retained twitching activity, although the proportion of motile cells varied substantially (22–99%). Notably, the partially phage-sensitive R30E and G75L mutants exhibited intermediate motile populations (26.9% and 43.2%, respectively) while maintaining twitching speeds comparable to those of several fully functional strains. These findings indicate that R30E and G75L support pilus assembly and force generation but compromise T4P function through a mechanism distinct from complete loss of pilus formation. We therefore hypothesized that these mutations alter the intrinsic mechanical properties of assembled pili and tested this hypothesis using single-molecule force spectroscopy.

### T4P stiffening compromises twitching motility

To directly link T4P structure, mechanics, and function, we quantified the force-dependent deformation of individual filaments using dual-trap optical tweezers. Purified PAO1 WT, PilA-R30E, and PilA-G75L pili were tethered between two optically trapped beads and subjected to controlled tensile loading while filament extension and restoring force were simultaneously monitored (**Fig. 4a**). Initial stretching frequently revealed multiple discrete rupture events, consistent with the presence of pilus bundles. Continued extension ultimately produced a final rupture event followed by force-free bead displacement, indicating that the remaining tether corresponded to a single T4P filament (Methods).

**Fig. 4:**
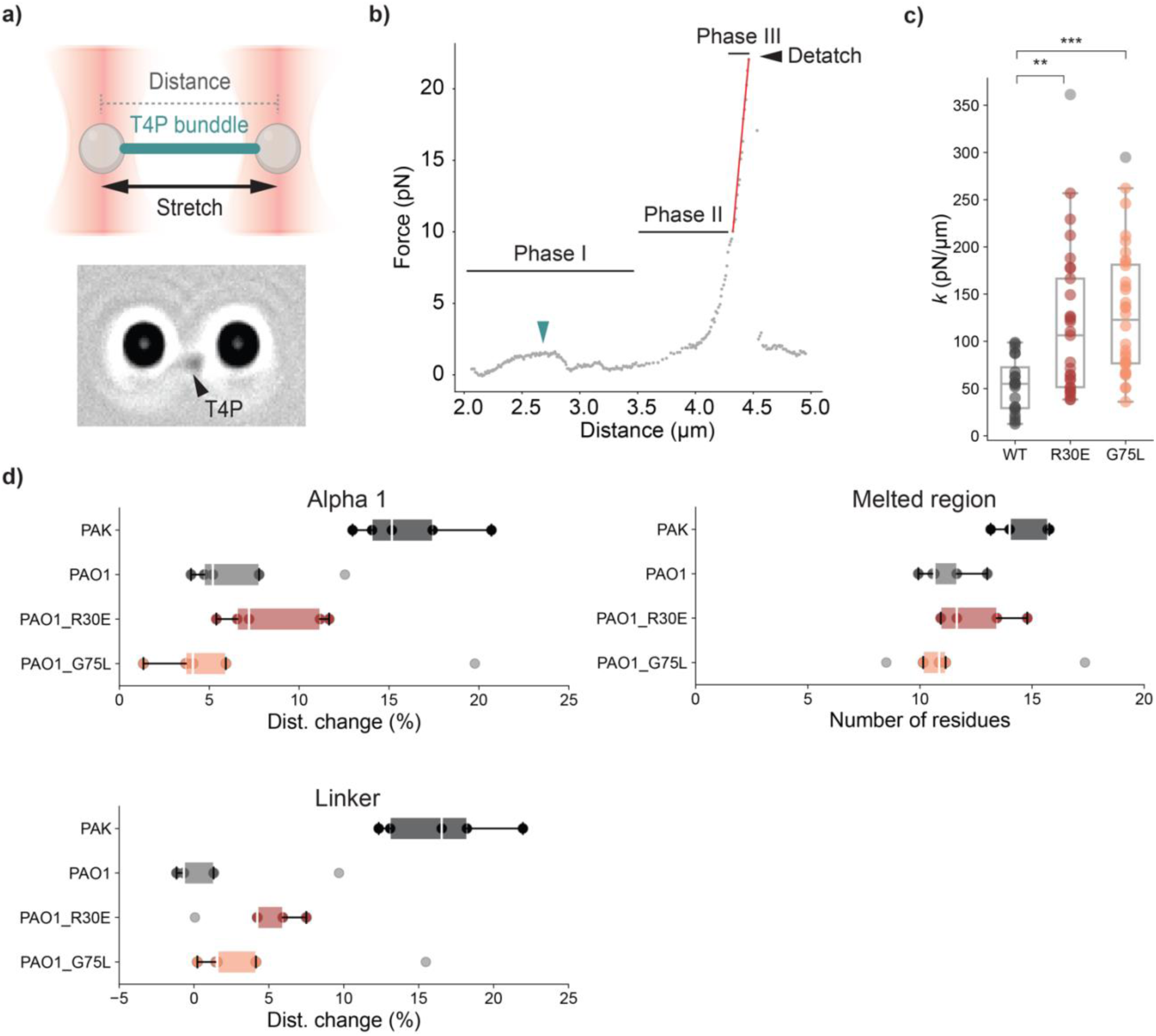
Single-molecule force spectroscopy reveals macroscopic stiffening of P. aeruginosa PAO1 T4P mutants. **a,** Schematic of the optical tweezers assay and a representative widefield view of the trapped bead–T4P complex. A purified T4P filament was tethered between two 2-µm carboxylate-modified polystyrene beads. The filament was stretched by displacing the right bead while the left bead is held stationary to record the force response. Inter-bead distance was measured between the centers of mass of the two beads. **b,** Representative force–distance curve of a single PAO1 T4P-WT. The curve illustrates distinct mechanical regimes: an initial resting state (Phase I), non-linear stretching (Phase II), and linear stretching (Phase III). The turquoise arrow indicates a minor initial force transient. The red line represents the calculated slope of the linear regime, from which the spring constant (k) was derived.19 **c,** Box plot comparing the macroscopic pilus stiffness (calculated spring constant, k) of T4P-WT and the T4P-R30E and T4P-G75L mutants. Each dot represents an independent pulling event; events failing quality control criteria (Methods) were excluded. The center line indicates the median, box limits represent the upper and lower quartiles, and whiskers indicate 1.5× the interquartile range (IQR). Statistical significance was assessed using a Kruskal-Wallis test followed by Dunn’s post-hoc test with Holm-Bonferroni correction for multiple comparisons (WT vs. R30E, P = 4.22 × 10-3; WT vs. G75L, 431 P = 4.80 × 10-5; R30E vs. G75L, P = 0.19). For simplicity in the plot legends in b, c, and d, WT and PAO1 refers to the PAO1 ΔfliC parental strain, and R30E and G75L mutants were generated in this ΔfliC background. Sample sizes are n = 20 (WT), 29 (R30E), and 32 (G75L) independent pulling events. **d,** Monomer extensibility of individual mutant pilins in silico. Box plots quantify the mechanical strain of the α1 helix (top), the mechanical strain of the linker region (middle), and force-induced deformation (number of melted residues) within the α1 hinge region (bottom) at the 100-ns snapshot of the steered molecular dynamics (SMD) simulations. Data represent the distribution across bulk subunits showed in Fig. 2a. Outliers are shown in gray. Data for T4P-WT of PAO1 and PAK are reproduced from Fig. 2d–f as a reference. n = 5 independent replicates per system

A representative force–extension curve obtained from a single WT pilus revealed four sequential mechanical regimes (**Fig. 4b**): a slack state (Phase I), a non-linear stretching regime (Phase II), a linear elastic regime (Phase III), and filament detachment. During the slack phase, bead displacement produced little force, indicating that the filament was not yet under tension. As extension increased, force rose non-linearly before transitioning to a linear regime characterized by a constant force–extension relationship. Final detachment was marked by an abrupt loss of tension, confirming rupture of the pilus tether. We interpret the non-linear regime as reflecting force-induced structural rearrangements within the filament, whereas the linear regime captures the intrinsic elasticity of the extended pilus and closely resembles the sustained loading conditions used in the SMD simulations. We therefore quantified filament stiffness by calculating the spring constant (*k*) from the slope of the linear regime.

Comparison of the spring constant (*k*) across multiple independent pulling events revealed that both mutant T4P variants were substantially stiffer than the WT T4P, with average values of 53.6 pN/µm (WT), 116.9 pN/µm (R30E), and 130.2 pN/µm (G75L) (**Fig. 4c**). Combined with the phage infection and motility phenotypes, these measurements demonstrate that R30E and G75L support T4P assembly but alter filament mechanics by increasing stiffness. Thus, disruption of the native interaction network does not weaken the filament; instead, it produces hyper-rigid pili with impaired biological function.

To determine how these mutations give rise to filament stiffening, we performed SMD simulations of the assembled R30E and G75L pili and analyzed their interaction networks under tension (**Extended Data Fig. 5**). The R30E substitution rewired the local electrostatic landscape, replacing the native R30–E49 interaction with a novel E30–R53 salt bridge and promoting force-stabilization of an E49–Y27 interaction. In contrast, G75L did not introduce major changes in the inter-subunit hydrogen-bonding or electrostatic networks. Despite these differences, neither mutant exhibited substantial changes in nanoscale pilin extensibility (**Fig. 4d**). Only modest alterations in α1 helix deformation, linker extension, and melted region unfolding were observed relative to WT PAO1. Together, these results indicate that the pronounced stiffening of R30E and G75L pili arises from subtle reorganization of the filament interaction network rather than large changes in the mechanical response of individual pilin subunits.

## Discussion

Our study reveals that the *P. aeruginosa* PAO1 T4P combine nanoscale rigidity with the macroscopic compliance required for biological activity. By comparative SMD analysis across six strains, we identify a conserved force-bearing network of inter-subunit interactions that governs force propagation through the filament and show that perturbing this network produces hyper-rigid pili that assemble normally yet exhibit impaired twitching motility and phage infectivity. Previous work has largely focused on the remarkable ability of T4P to withstand the extraordinary forces generated during retraction^25,42,43^. Our findings suggest that the relevant evolutionary constraint is not maximal mechanical stability, but the ability to balance force resistance with filament dynamics. We propose that T4P operate within a narrow biophysical regime in which inter-subunit interactions must be sufficiently strong to sustain force transmission while remaining sufficiently labile to permit pilin extraction during retraction and subsequent reassembly.

Structural analyses of T4P have catalogued extensive networks of inter-subunit contacts, yet their functional role under tension has remained largely unknown. More broadly, a recurring challenge in mechanobiology is distinguishing static structural contacts from those that actively bear and transmit force. Similar principles are observed in cytoskeletal polymers, where specific inter-subunit interfaces and structural states coordinate to govern filament mechanics and dynamics^44^. Here, by combining SMD simulations with single-molecule force spectroscopy, we identify a conserved subset of T4P interactions that persist under tension and function as force-bearing elements within the filament. Identifying these mechanically active contacts provides a framework for understanding how specific molecular bonds tune the emergent mechanics of dynamic assemblies.

A striking observation was the disconnect between local and global behavior. Although the R30E and G75L mutations produced pronounced filament stiffening in optical tweezer measurements, they caused only modest changes in the deformation of individual pilin subunits during SMD simulations. This suggests that T4P flexibility emerges from collective interactions distributed across the filament rather than from the intrinsic extensibility of individual pilins. This trade-off between local subunit rigidity and global filament flexibility highlights a central challenge in molecular biomechanics: the macroscopic behavior of supramolecular assemblies often cannot be inferred directly from local structural changes. Bridging this gap will require integrative approaches capable of connecting atomistic interactions to global mechanical properties across multiple length scales.

## Author contributions

Conceptualization: C.T., A.P, and Y.K.

Data curation: C.T. and Y.K.

Formal analysis: C.T. and Y.K.

Funding acquisition: C.T., A.P, and Y.K.

Investigation: C.T. and Y.K.

Methodology: C.T., L.L.B., F.E, Y.K., and M.N.

Resources: Z.A.

Supervision: A.P. and Y.K.

Visualization: C.T.

Writing-original draft: C.T., A.P., and Y.K.

Writing-review and editing: C.T., A.P., Y.K., and M.N.

## Acknowledgements

We acknowledge Florence Pojer, Yoan Duhoo, and Kelvin Lau from the EPFL Protein Production and Structure Core Facility for their advice and assistance with cryo-EM grid preparation and data analysis. We thank the team at the Dubochet Center for Imaging (DCI) in Lausanne for their support with the collection and processing of electron microscopy data. We thank Aleksandar Antanasijevic at EPFL for his guidance on initial cryo-EM data processing, analysis, and model building. We also thank Rob Lavigne at KU Leuven for kindly providing the LUZ19 phage, and Olivera Francetic at Institut Pasteur for her valuable feedback during the preparation of this manuscript.

---

**Extended Data Fig. 1:**
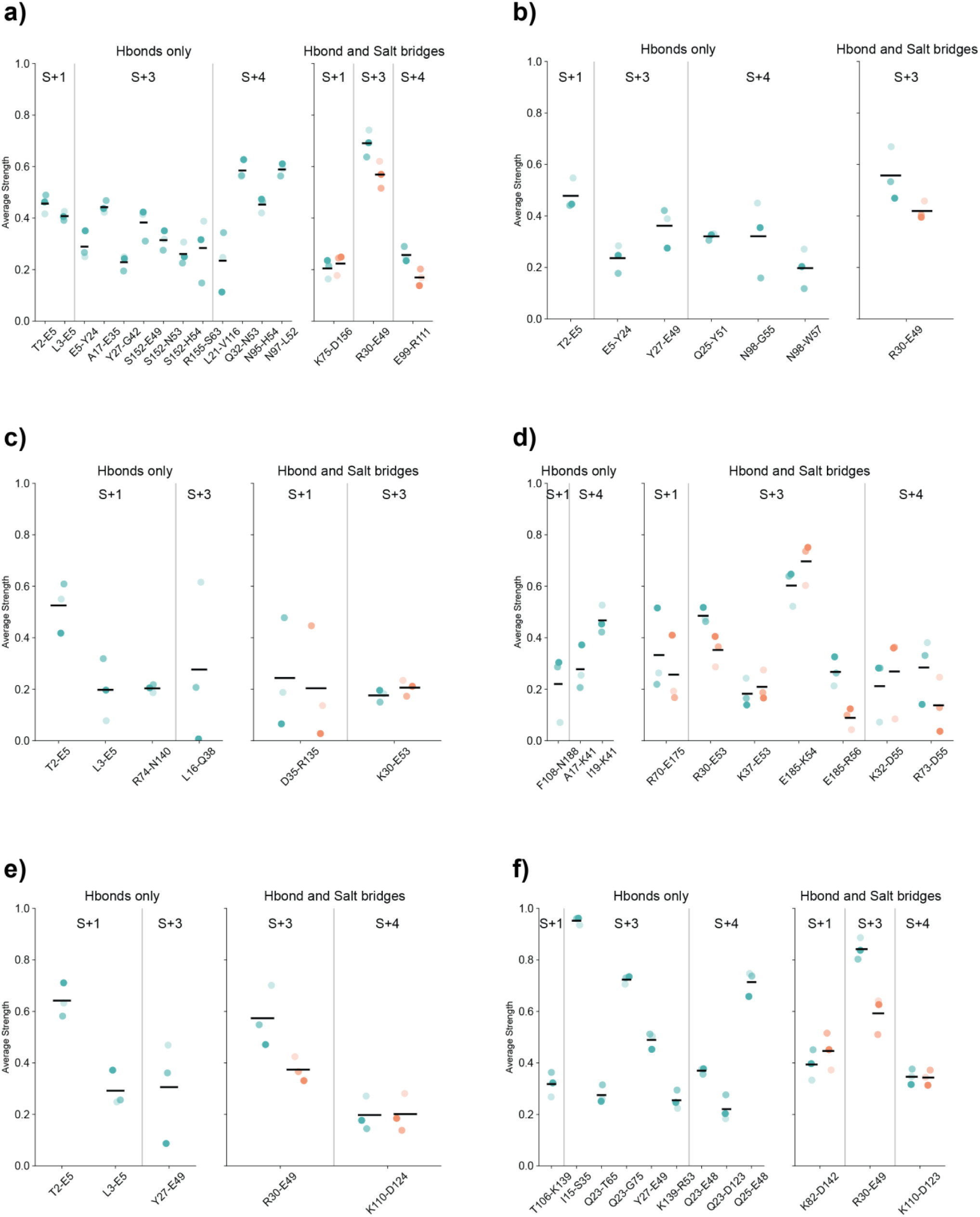
Inter-subunit interaction networks from equilibrium molecular dynamics (MD) simulations across six bacterial strains. a–f, Interaction networks for *N. meningitidis* (*Nm*) (a), *N. gonorrhoeae* (*Ng*) (b), *E. coli* (*Ec*) (c), *M. xanthus* (*Mx*) (d), and *P. aeruginosa s*trains PAK (e) and PAO1 (f). In each panel, the left plot displays networks characterized exclusively by hydrogen bonds (Hbonds, turquoise), and the right plot displays networks involving both hydrogen bonds and salt bridges (salmon). Each circle represents the mean contact persistence from a single equilibrium MD replicate. Horizontal black lines indicate the overall mean (*n* = 3 independent replicates).

**Extended Data Fig. 2:**
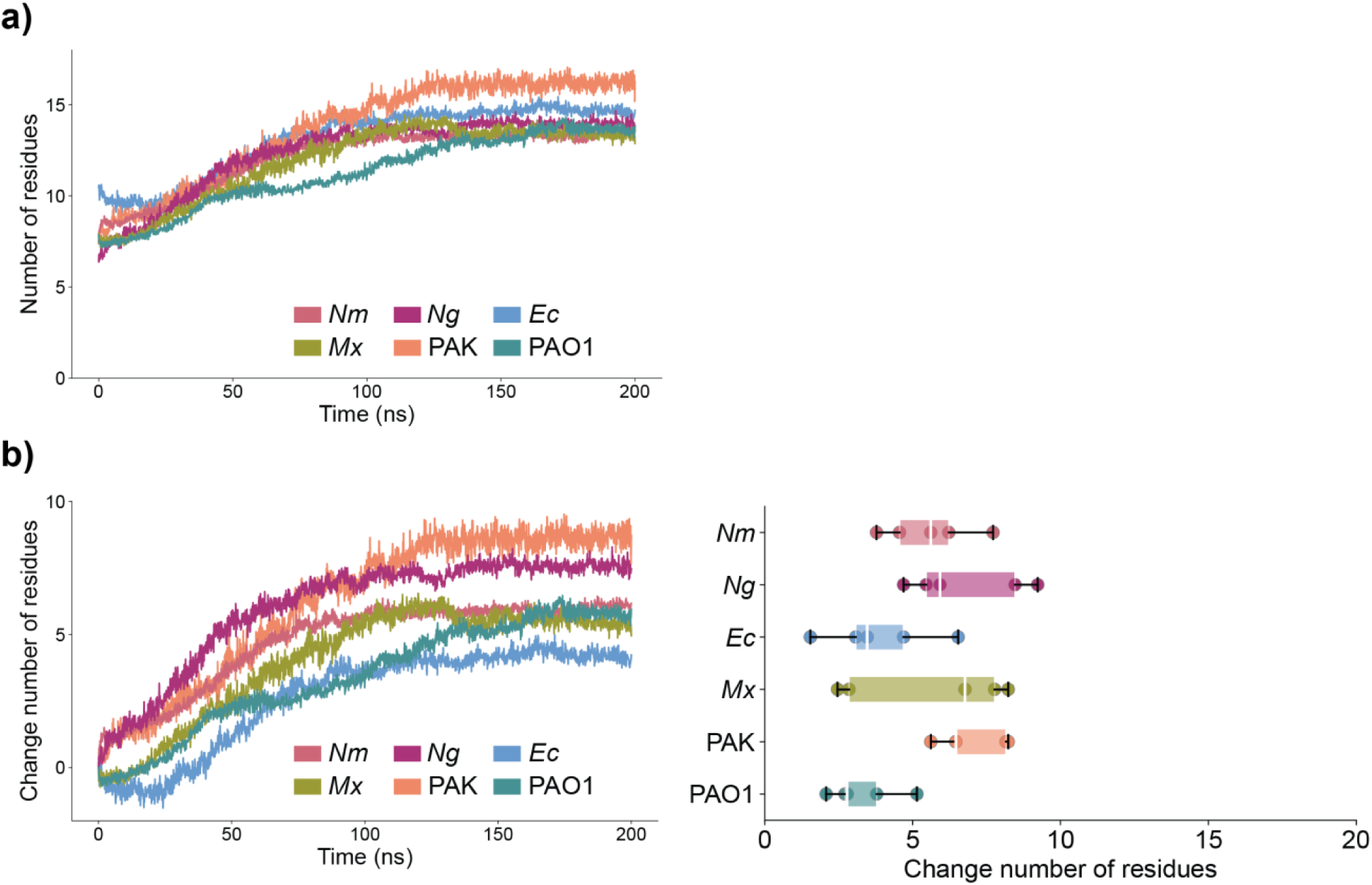
Unfolding dynamics of the α1 helix melted regions. **a,** Time evolution of the absolute number of unstructured (non-α-helical) residues in the melted segments, determined by DSSP analysis, for *N. meningitidis* (*Nm*), *N. gonorrhoeae* (*Ng*), *E. coli* (*Ec*), *M. xanthus* (*Mx*), and *P. aeruginosa* strains PAK and PAO1. These curves represent the full 200-ns evolution corresponding to the 120-ns time windows shown in Fig. 2f. **b,** Left: Time evolution of the net change in the number of these unstructured residues. Right: Distribution of melted residues at 100 ns across bulk subunits for each strain; *n* = 5 replicates. For box plots, the center line indicates the median, box limits represent the upper and lower quartiles, and whiskers indicate 1.5× the interquartile range (IQR).

**Extended Data Fig. 3:**
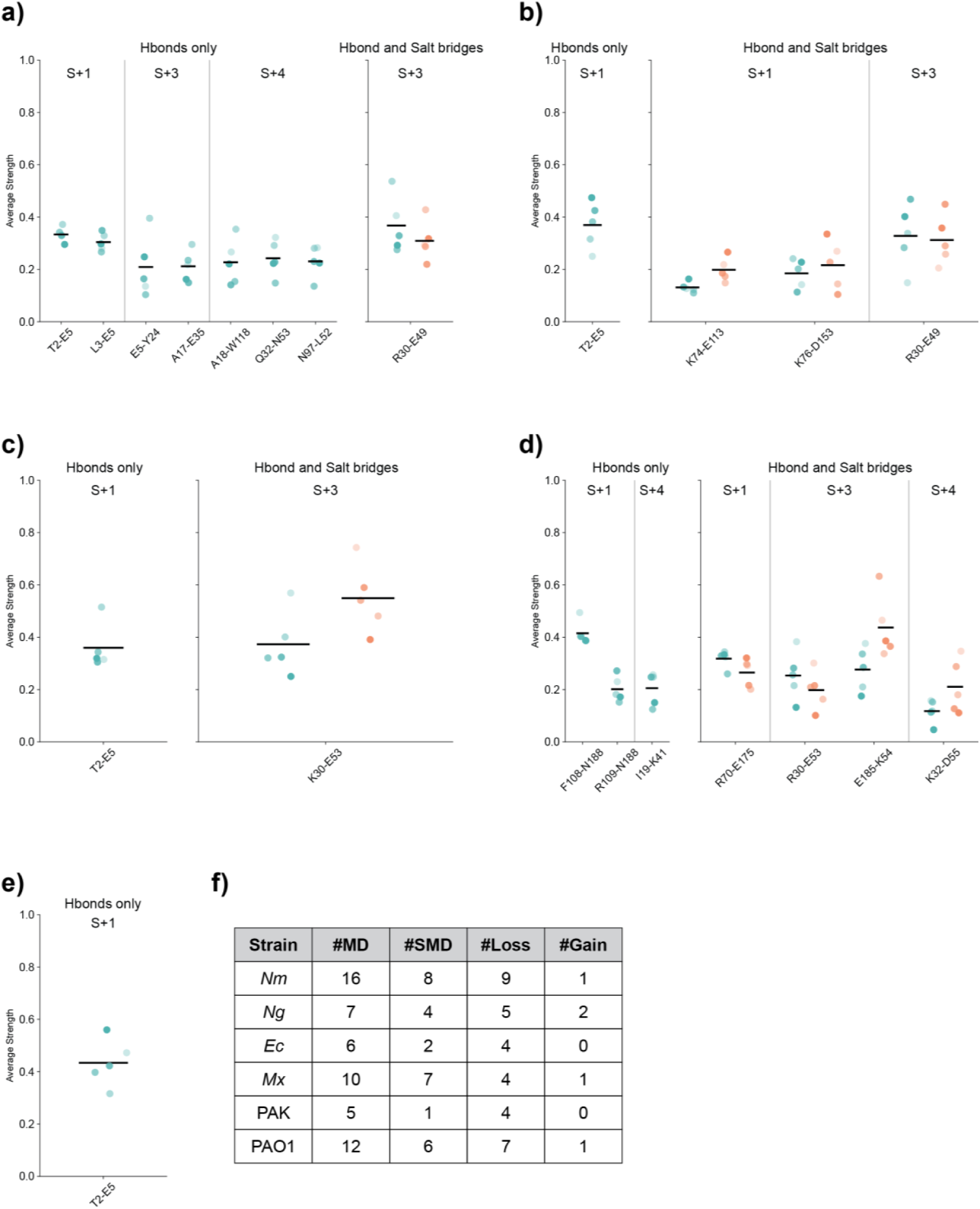
Inter-subunit interaction networks from steered molecular dynamics (SMD) simulations across five bacterial strains. a–e, Interaction networks for *N. meningitidis* (*Nm*) (a), *N. gonorrhoeae* (*Ng*) (b), *E. coli* (*Ec)* (c), *M. xanthus* (*Mx*) (d), and *P. aeruginosa* strain PAK (e). In each plot, the left panel displays networks characterized exclusively by hydrogen bonds (Hbonds, turquoise), and the right panel displays networks involving both hydrogen bonds and salt bridges (salmon). Each circle represents the mean contact persistence from a single SMD replicate. Horizontal black lines indicate the overall mean (*n* = 5 independent replicates). **f,** Summary of inter-subunit interactions during equilibrium MD and SMD. #MD: number of interactions identified in the equilibrium MD simulations. #SMD: number of interactions identified in the SMD simulations. #Loss: number of interactions lost during SMD compared to MD. #Gain: number of novel interactions gained during SMD compared to MD.

**Extended Data Fig. 4:**
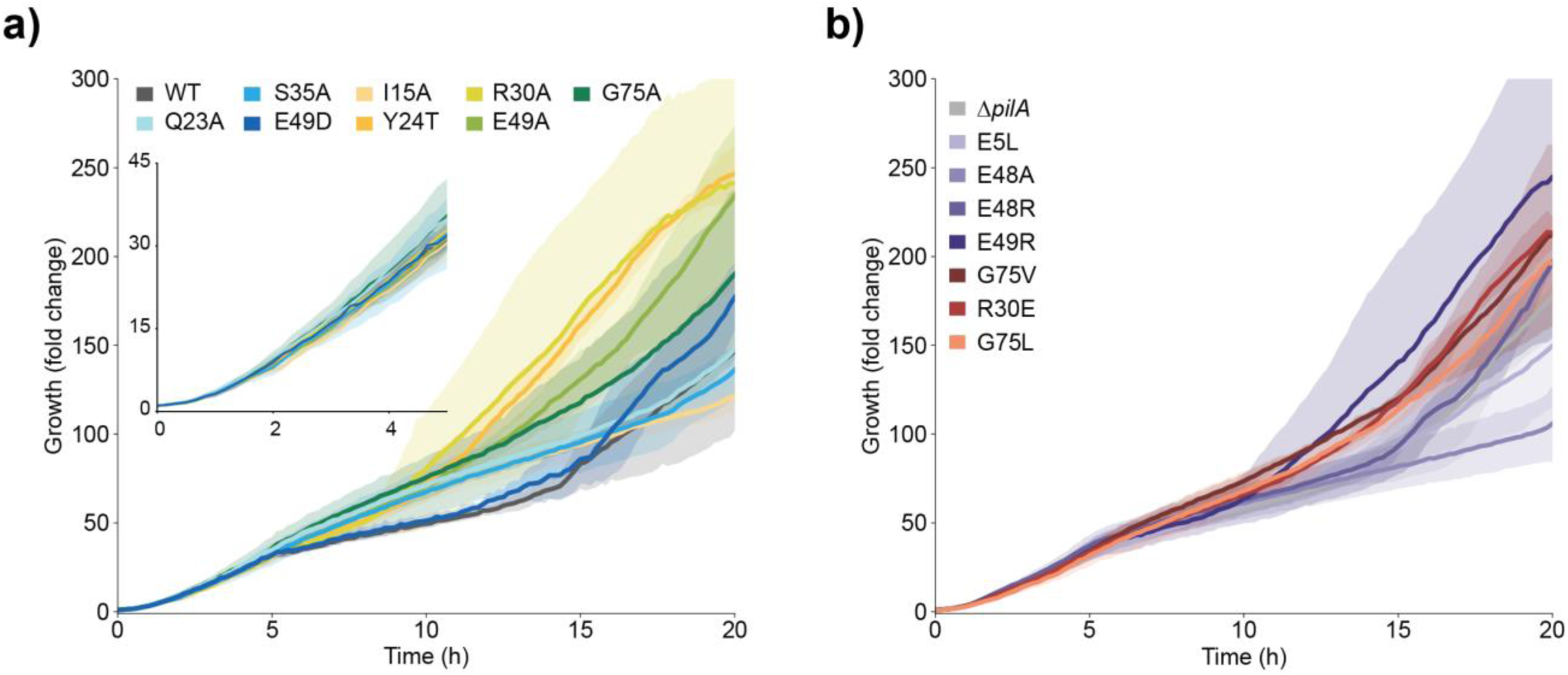
Growth curves of *P. aeruginosa* PAO1 PilA-WT, Δ*pilA*, and PilA mutants. Bacterial density (fold-change relative to initial concentration) was monitored over 20 h in the absence of phage. **a, b,** Growth controls for the phage-sensitive strains corresponding to the experiments in Fig. 3b (a), and the phage-resistant and partially sensitive strains corresponding to the experiments in Fig. 3c (b). All experiments were performed in independent biological triplicates (*n* = 3).

**Extended Data Fig. 5:**
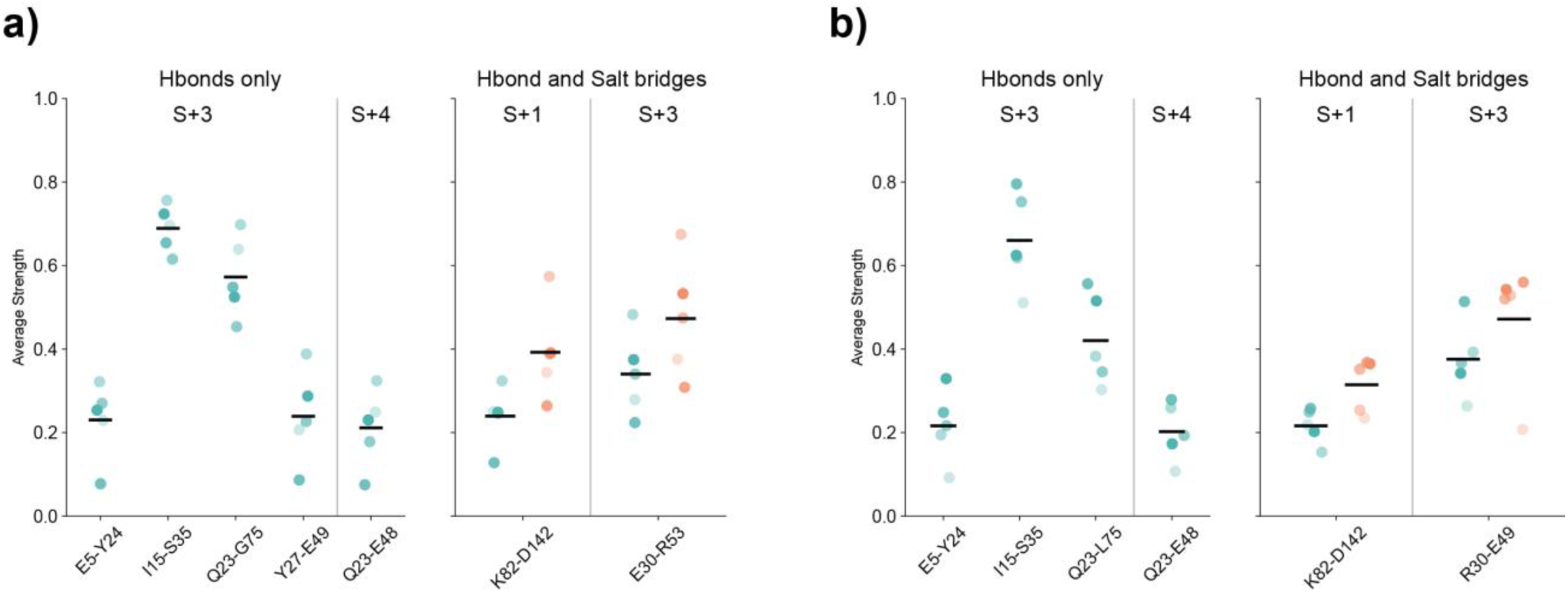
Inter-subunit interaction networks from steered molecular dynamics simulations of *P. aeruginosa* PAO1 T4P-R30E and T4P-G75L. a, b, Interaction networks for the T4P-R30E (a) and T4P-G75L (b) mutants. In each panel, the left plot displays networks characterized exclusively by hydrogen bonds (turquoise), and the right plot displays networks involving both hydrogen bonds and salt bridges (salmon). Each circle represents the mean contact persistence from a single SMD replicate. Horizontal black lines indicate the overall mean (*n* = 5 independent replicates).

## Extended Tables

**Extended Data Table 1.** The list of investigated T4P in this study. The structural details of the studied systems are reported in this table.

| Bacterial strain | PDB code | Resolution (Å) | #Subunits | #Residues |
| --- | --- | --- | --- | --- |
| *Neisseria meningitidis* (Nm) | 5KUA | 6 | 26 | 161 |
| *Neisseria gonorrhoeae* (Ng) | 5VXX | 5.1 | 21 | 156 |
| *Enterohemorrhagic*<br>*Escherichia coli* (Ec) | 6GV9 | 8 | 14 | 140 |
| *Myxococcus xanthus* (Mx) | 8TJ2 | 3 | 21 | 208 |
| *Pseudomonas aeruginosa* PAK | 5VXY | 8 | 21 | 144 |
| *Pseudomonas aeruginosa* PAO1 | 32AY | 2.8 | 22 | 143 |

**Extended Data Table 2.** The list of performed classical and Steered MD simulations. All studied systems, as well as number of replicates and simulation time are reported here.

| Simulation | T4P name | Simulation length (ns) | #Replicates | #Atoms | #Subunits | Box size (Å<sup>3</sup>) |
| --- | --- | --- | --- | --- | --- | --- |
| **MD** | Nm | 3000 | 3 | 357,327 | 26 | 100*100*367 |
|  | Ng | 3000 | 3 | 298,856 | 21 | 100*100*310 |
|  | Mx | 3000 | 3 | 297,709 | 21 | 100*100*312 |
|  | PAK | 3000 | 3 | 309,783 | 21 | 100*100*320 |
|  | PAO1 | 3000 | 3 | 313,714 | 22 | 100*100*329 |
| **SMD** | Nm | 200 | 5 | 621,927 | 26 | 102*102*600 |
|  | Ng | 200 | 5 | 595,775 | 21 | 100*100*600 |
|  | *Ec* | 200 | 5 | 692,554 | 21 | 100*100*700 |
|  | *Mx* | 200 | 5 | 633,563 | 21 | 100*100*640 |
|  | PAK | 200 | 5 | 482,236 | 21 | 90*90*600 |
|  | PAO1 | 200 | 5 | 508,081 | 22 | 89*89*650 |

## Materials and Methods

### Studied systems

We studied the dynamics of six different T4P: the *N. meningitidis* (*Nm*), *N. gonorrhoeae* (*Ng*), *E. coli* EHEC (*Ec*), *M. xanthus* (*Mx*), *P. aeruginosa* PAK and PAO1. The 3D coordinates of the studied systems were retrieved from the Protein Data Bank^45^ and the details are reported in **Extended Data Table 1**. For the MD and SMD simulations of *Ec*, the number of subunits was increased from 14 to 18 and 21, respectively, by applying the internal helical symmetry parameters. We refer to bulk subunits as the central subunits in the pilus, after excluding 4 subunits from the top and 4 from the bottom of the pilus. All the analyses in this study were performed on the bulk subunit, unless stated otherwise. The environment of the histidines was manually checked and consequently protonated with a hydrogen at the ɛ nitrogen (H54 and H149 for *Nm*, H54 and H146 for *Ng, and* H54 for *Ec*).

### Molecular dynamics simulations

#### System preparation

The MD simulations of T4P were performed in a POPE (1-palmitoyl-2-oleoyl-sn-glycero-3-phosphoethanolamine) model membrane. The T4P were initially placed near the membrane with the linker and N-terminus of the first subunit partially inserted into the bilayer. All systems were prepared using CHARMM-GUI membrane builder server^46,47^ and the CHARMM36m force-field parameter set^48,49^: *(i)* the conformation of the linker in the α1 helix of the membrane-embedded bottom subunit was modeled as a continuous helix using Modeller software, *(ii)* hydrogen atoms were added, *(iii)* the POPE model membrane was considered to build the inner bacterial membrane, *(iv)* the solute was hydrated with a rectangular box of explicit TIP3P water molecules with a buffering distance up to 14 Å, *(v)* Na^+^ and Cl*^−^* counter-ions were added to reproduce a physiological salt concentration (100 mM solution of sodium chloride). For each system, three replicates of 3-*µ*s MD simulations were performed. Details of all the simulations are reported in **Extended Data Table 2**.

#### Production of the trajectories

The GROMACS 2020.4 package was used to carry out all simulations^50^. The energy minimization was performed using a steepest descent (SD) algorithm for 10000 steps, to minimize any steric overlap between system components. This was followed by an equilibration simulation in an NPT ensemble at 310 K, allowing the lipid and solvent components to relax around the restrained protein. All the protein and lipid non hydrogen atoms were harmonically restrained, with the constraints gradually reduced in 6 distinct steps with a total of 0.375 ns. The particle mesh Ewald algorithm^51^ was applied to calculate electrostatic forces, and the van der Waals interactions were smoothly switched off at 10-12 Å by a force-switching function^52^. A time step of 1 fs was used in all simulations. The productions were realized in the NPT ensemble. The time step was set to 2.0 fs. The temperature was kept at 310 K, and pressure at 1 bar using the Langevin piston coupling algorithm. The SHAKE algorithm was used to freeze bonds involving hydrogen atoms, allowing for an integration time step of 2.0 fs. The Particle Mesh Ewald (PME) method^53^ was employed to treat long-range electrostatics. Half-harmonic potentials were applied at the tip of the pilus (subunit A-D), in order to prevent the dissociation of the tip. PLUMED 2.8.0^54^ and the PLUMED-ISDB^55^ module were used to add lower and upper walls on the distance between the *Cα* atoms of four pairs of residues (*Ec*: *T*45*_A_*-*L*52*_B_*, *T*45*_B_*-*L*52*_C_*, *T*45*_C_*-*L*52*_D_*, *L*49*_D_*-*K*30*_A_*, PAK: *T*45*_A_*-*S*52*_B_*, *T*45*_B_*-*S*52*_C_*, *T*45*_C_*-*S*52*_D_*, *E*49*_D_*-*R*30*_A_*, *Nm*: *S*45*_A_*-*L*52*_B_*, *S*45*_B_*-*L*52*_C_*, *S*45*_C_*-*L*52*_D_*, *E*49*_D_*-*R*30*_A_*, *Ng*: *S*45*_A_*-*L*52*_B_*, *S*45*_B_*-*L*52*_C_*, *S*45*_C_*-*L*52*_D_*, *E*49*_D_*-*R*30*_A_*, *Mx*: *A*46*_A_*-*S*52*_B_*, *A*46*_B_*-*S*52*_C_*, *A*46*_C_*-*S*52*_D_*, *S*49*_D_*-*R*30*_A_*, PAO1: *T*45*_A_*-*S*52*_B_*, *T*45*_B_*-*S*52*_C_*, *T*45*_C_*-*S*52*_D_*, *E*49*_D_*-*R*30*_A_*) with a force constant of 1000 kcal/mol/U^2^. The choice of residues has been made according to their fluctuations, and those with the lowest deviations were selected. The coordinates of the system were written every 100 ps. For every system, three replicates were performed, starting with different initial velocities. MD simulations of the *Ec* T4P were taken from our previous study^29^, which consisted of three replicates of 3 *µ*s in the presence of calcium ions and performed in the same conditions as the present study and using GROMACS 2019.4 and PLUMED 2.6.0.

#### Steered molecular dynamics simulations

For the steered molecular dynamics (SMD) simulations, the systems were prepared using the CHARMM36m force-field parameter set^48,49^: *(i)* the 3D coordinates were extracted from the PDB (as reported in **Extended Data Table 1** and in the case of *Ec* the number of subunits were increased to 21, *(ii)* hydrogen atoms were added, *(iii)* the solute was hydrated with a triclinic box of explicit TIP3P water molecules with a buffering distance up to 14 Å in the direction of x- and y-axis and a larger distance between 600 to 700 Å for the z-axis, *(iv)* Na^+^ and Cl^-^counter-ions were added to reproduce a physiological salt concentration of 100 mM. The details of the box size are reported in **Extended Data Table 2**. The SMDs were carried out at constant velocity with the spring constant of 1000 kJ/(mol·nm²) and pulling velocity of 1 Å/ns. For each system, we performed five replicates of 200 ns, and the coordinates of each simulation were written every 10 ps. The force was applied directly to the atoms in the top four subunits of T4P and the atoms in the first 30 residues of the four bottom-most subunits were fixed.

### Structural analysis

#### Stability of the trajectories

Standard analyses of the MD trajectories were performed using the gmx module of GROMACS 2020.4. The root mean square deviation (RMSD) of backbone atoms (*Cα, C, N, O*) from the initial frame were recorded along each replicate (**Supplementary Data Fig. 1**). Based on the RMSD profiles, we performed the subsequent analysis over the subset of simulations where the systems are fully relaxed, *i.e.* considering the last 2500 ns for the classical MD simulations. The by-residue root mean square fluctuations (RMSF) was computed over the backbone atoms (*Cα, C, N, O*), with respect to the average conformation (**Supplementary Data Fig. 2**). All the studied systems remained stable along the MD trajectories.

#### Distances and lengths

The change in length of the α1 helix was measured as the distance between the *Cα* atoms of residues 1 and 53 throughout the simulations, normalized to the initial α1 length from the respective cryo-EM structure. Similarly, the change in linker length was defined as the distance between the corresponding *Cα* atoms (G14–P22 for *Nm, Ng, Mx*, PAK, and PAO1; G11–P22 for *Ec*), normalized to the initial cryo-EM linker length. The melted region was quantified by counting the number of unstructured residues not assigned as an α-helix (H) by DSSP^40^. The initial melted regions for each system were defined as G14–P22 for *Nm, Ng*, and PAK; G11–P22 for *Ec*; and I19–N23 for *Mx* and PAO1. The net change in the melted region was calculated by subtracting the initial number of non-helical residues from the number observed at each simulation time step. To eliminate edge effects, these analyses were performed exclusively on the “bulk” subunits, defined as the intermediate set of monomers remaining after excluding the top four and bottom four subunits of the filament.

#### Salt bridges and hydrogen bonds

We used VMD to record the average distances between pairs of residues that form salt-bridges along the simulation and merged the results from all the replicates of each system by collecting the values of every pair. This distance is measured between the center of mass of the oxygens in the acidic side chains and center of mass of the nitrogens in the basic side chains. For a given salt bridge between residues *i* and *j*, an interaction strength is computed as the percentage of conformations in which the measured distance is at most 4 Å between residues *i* and *j*. We identified hydrogen-bonds (Hbonds) using the HBPLUS algorithm^56^. Hbonds are detected between donor (D) and acceptor (A) atoms that satisfy the following geometric criteria: *(i)* maximum distances of 3.9 Å for D-A and 2.5 Å for H-A, *(ii)* minimum value of 90*^◦^* for D-H-A, H-A-AA and D-A-AA angles, where AA is the acceptor antecedent. For a given Hbond between residues *i* and *j*, an interaction strength is computed as the percentage of conformations in which the HBond is formed between any atoms of the same pair of residues (*i* and *j*). To account for the helical symmetry of T4P, we merged the inter subunit interactions from all the replicates of the bulk subunit. For a given pair of inter subunit interaction (either a salt bridge or an Hbond), we reported all strength values, if the interaction happens at least between three subunits with the minimum stability of 50% (0.5) and has an average stability of at least 20% (calculated over all the subunits and all the replicates).

### Computational tools

Trajectories generated by MD simulations were analyzed using gmx rms, gmx rmsf, gmx sasa utilities of GROMACS 2019.4 4^50^. VMD^57^ and PyMOL^58^ were used for visualization and plots were generated using the R software^59^, and Python^60^.

### Bacterial strains and growth conditions

*Pseudomonas aeruginosa* PAO1 strains, all generated in a flagellum-deficient *ΔfliC* background, were used for pili purification, cryo-EM, phage infection, single-cell twitching motility, and optical tweezer experiments. *E. coli* strains XL10-Gold and S17.1 were used for plasmid construction and conjugative mating, respectively. Liquid cultures of all strains were grown in LB medium at 37 °C with shaking at 280 rpm. Solid LB agar (1.5% wt/vol agar) for pili purification was prepared by the Infrastructures Unit of the School of Life Sciences, EPFL. Gentamicin was added for selection at 10 µg/mL for *E. coli* and 60 µg/mL for *P. aeruginosa*.

### Strain and plasmid construction

Strains and plasmids, together with their corresponding oligonucleotides, are listed in **Supplementary Data Table 1, 2, and 3**. All gene deletions and insertions were performed by two-step allelic exchange using the *sacB*-based suicide vector pEX18Gent^61^. For in-frame deletions, ∼500-1,000 bp regions immediately upstream and downstream of each target locus were PCR-amplified with Phusion High-Fidelity DNA Polymerase (Thermo Fisher Scientific, #F534S), with primer design preserving three native codons at both the 5’ and 3’ boundaries. Point substitutions were introduced by site-directed mutagenesis using complementary mutagenic primers. PCR products and EcoRI/XbaI-linearized pEX18Gent were assembled by Gibson assembly^62^ and transformed into *E. coli* XL10-Gold. Plasmids were then introduced into *E. coli* S17.1 for conjugative mating into *P. aeruginosa*. In-frame insertion of full-length *pilA* alleles bearing the specified point mutations was carried out in a *ΔfliCΔpilA* double-mutant background using the same exchange workflow. Recombinant *P. aeruginosa* strains were confirmed by colony PCR (Promega, #M7823) and Sanger sequencing.

### Purification of T4P from *P. aeruginosa ΔfliC* and *pilA* mutants

Pili were purified from the flagellum-deficient *P. aeruginosa* PAO1 *ΔfliC* strain to avoid flagellum contamination, using a protocol modified from previous^17,35^. Briefly, 50 µL of an overnight LB culture was spread onto LB agar plates and incubated at 30 °C for 24 h. Bacterial lawns from ten plates were scraped and resuspended in 8 mL of Buffer 1 (150 mM ethanolamine, pH 10.5, Sigma, #15014, supplemented with 1 mM DTT, Fluorochem, #M02712). Pili were sheared by three cycles of vigorous vortexing (1 min) followed by chilling on ice (1 min). Cells and cellular debris were removed by four successive centrifugations at 8,000g for 30 min at 4 °C. The combined supernatants were adjusted to 10% ammonium sulfate (Sigma-Aldrich, #A4915) saturation and incubated at 16 °C for 16 h. Precipitated pili were collected by centrifugation at 13,000g for 60 min at 4 °C. Pellets were then resuspended in 8 mL of Buffer 2 (20 mM ethanolamine, pH 10.5) and subjected to a second 10% ammonium sulfate precipitation at ambient temperature for 24 h. The final pellet was resuspended in 500 µL of Buffer 2 containing 1 mM DTT and stored at -20 °C. For optical tweezer experiments, pili bearing specific point mutations were purified from a hyperpiliated PAO1 *ΔfliCΔpilTΔpilH* background strain using the identical workflow.

### Cryo-EM sample preparation and data collection

A 3-µL aliquot of the 0.2 mg/mL purified T4P sample was applied onto the Quantifoil Cu 1.2/1.3 400 mesh grid (Thermo Fisher Scientific), glow discharged with GloQube for 90 s and 15 mA current (Quorum Technologies). The grids were blotted for 4 s at 10 °C and 95% humidity (blot force 4) and subsequently plunge-frozen in liquid ethane using a Vitrobot Mark IV system (Thermo Fisher Scientific). Grids were imaged on an X-FEG fringe-free 200-kV Glacios transmission electron microscope (Dubochet Center for Imaging, Lausanne, Switzerland) equipped with a Falcon 4 direct electron detector (Thermo Fisher Scientific) operating in electron counting mode. Automated data collection was controlled using EPU v.3.10 software utilizing aberration-free image shift (AFIS). Images were collected in fast mode at a nominal magnification of 150,000×. While the nominal pixel size at the specimen level was 0.926 Å, the calibrated pixel size used for processing was 0.95 Å. For helical reconstruction of the T4P, movies comprising 40 frames were collected over a nominal defocus range of -0.8 to -2.6 µm, with a total accumulated dose of 40 e^-^/Å^2^, yielding a total of 4,224 movies stored as electron event recording (EER) files.

### Cryo-EM image processing and 3D reconstruction

All cryo-EM image processing and helical reconstructions were performed in CryoSPARC (v.4.1)^63^. Movies were initially imported into the Patch Motion Correction job for dose-weighted, anisotropic motion correction, and subsequently processed with Patch CTF Estimation to fit per-micrograph defocus and astigmatism. Micrographs were curated using the Manually Curate Exposures job with thresholds of CTF fit resolution < 5 Å, average defocus < 30,000 Å, and astigmatism < 2,000 Å, resulting in 2,215 high-quality micrographs. Filament segments were extracted using the Filament Tracer job, with diameters set to 30 Å and 100 Å based on the known pilus width of ∼52 Å. A total of 281,910 filament segments were extracted in 400-pixel boxes, and 2D classification yielded 157,159 particles from well-resolved classes for 3D reconstruction. An average power spectrum of a representative 2D-class segment was analyzed using the Helixplorer-1 online server^64^ to estimate initial helical symmetry, yielding a rise of 10.191 Å and a twist of 88.467 degrees. These parameters guided the initial Helical Refinement, which enforces strict global symmetry, generating a 3.07 Å map. Because strict helical symmetry constraints can blur flexible regions that deviate from perfect symmetry (such as loops, inter-subunit interfaces, or flexible termini), we subsequently applied non-uniform refinement to the helical map. Non-uniform refinement retains global symmetry but introduces adaptive regularization that down-weights poorly ordered regions while reinforcing the well-aligned core. This targeted approach enhanced the signal-to-noise ratio in the most rigid regions of the filament while preserving high-resolution features in flexible areas, ultimately improving the final map resolution to 2.76 Å, as determined by the gold-standard Fourier shell correlation (FSC) of 0.143.

### Model building and refinement

The atomic model of the homologous *P. aeruginosa* PAK T4P major pilin (PDB: 5XVY)^17^ was used as an initial template. This monomer was rigid-body fitted into the sharpened 2.8 Å *P. aeruginosa* PAO1 cryo-EM density map using UCSF ChimeraX v1.17.3^65^. The sequence was mutated to match PAO1 PilA, and the model was manually rebuilt in Coot v.0.9.8.7^66^ to accurately position side chains into the high-resolution density. The single monomer then underwent an initial real-space refinement against the sharpened map using Phenix v.1.20.1-4487^67^. To reconstruct the full helical filament, we utilized the built-in symmetry tools in ChimeraX. The measure symmetry function was applied to the cryo-EM map to determine the helical rise and twist angle per subunit, establishing the central helical axis. These measured symmetry parameters were then applied to generate the full helical assembly from the fitted monomer. The monomer placement was finely adjusted and symmetry-expanded until the repeated subunits tracked the experimental density map. Finally, the assembled multi-subunit pilus was manually modified in Coot to optimize inter-subunit interfaces and fit the local density. This complete assembly was then refined in Phenix. This cycle of manual rebuilding in Coot and real-space refinement in Phenix was repeated iteratively until the overall geometry, inter-subunit contacts, and model-to-map fit scores were optimal.

### Phage infection assay

*P. aeruginosa* PAO1 *ΔfliC* background wild-type, *ΔpilA*, and strains harboring the indicated *pilA* mutations were inoculated in 1 mL of Luria-Bertani (LB) broth and incubated overnight at 37 °C. The overnight cultures were diluted 1:200 into 2 mL of fresh LB and grown at 37 °C for 2 hours to reach the exponential phase, after which the optical density at 600 nm (OD_600_) was adjusted to approximately 0.3 (corresponding to ∼10^8^ CFU/mL). LUZ19 bacteriophage (a gift from Rob Lavigne, KU Leuven) stored at 4 °C in LB was diluted to achieve a multiplicity of infection (MOI) of 10^-4^. For the infection assay, 10 µL of the adjusted bacterial culture was mixed with 290 µL of the diluted LUZ19 phage^41^ to a final volume of 300 µL. This mixture was immediately transferred into a 96-well plate (Greiner Bio-One, #655180), with three technical replicates per strain. Bacterial growth (OD_600_) was monitored continuously using a plate reader (SPECTROstar Nano, BMG LABTECH) at 37 °C. Measurements were taken every 10 minutes for 121 cycles (20 hours), preceded by 10 seconds of shaking at 300 rpm. Growth curves were generated by subtracting the LB blank background and normalizing the values to the initial OD_600_ reading (t = 0). Data processing and plotting were performed using custom Python scripts (version 3.9.16).

### Single cell twitching motility assay

Bacterial cells were inoculated and incubated overnight at 37 °C in LB. Cultures were then diluted to an OD_600_ of 0.02 and grown to mid-exponential phase (OD600 ∼0.2-0.5), before being adjusted to an OD_600_ of 0.2. Tryptone agarose plates (0.5% tryptone, 0.25% NaCl, and 0.5% agarose) were prewarmed at ambient temperature for 45-60 minutes. A 14-mm diameter tryptone agarose pad was excised, and a 1 µL droplet of the adjusted bacterial culture was deposited onto the top surface. The droplet was allowed to absorb for 30 seconds before the pad was inverted and placed onto a glass-bottom dish (MatTek, P35G-1.5-20-C), ensuring the cells were in direct contact with the glass. Samples were incubated at 37 °C for 2 hours prior to imaging. Time-lapse imaging was performed at ambient temperature using a Nikon ECLIPSE Ti2 inverted microscope equipped with a Photometrics Prime 95B camera and controlled by NIS-Elements software (version AR 5.41.01). Phase-contrast images were acquired using a 40x Plan Apo phase-contrast objective (NA 0.9). To capture twitching motility, image sequences were recorded at 2-second intervals for a total of 5 minutes. Cell segmentation and annotation were performed using the deep neural network image-segmentation algorithm Omnipose^68^. The ImageJ plugin TrackMate^69^ was then used for cell tracking, with tracking errors subjected to manual correction. Motile cells were defined by a minimum displacement threshold of 0.08 µm per frame over at least 3 consecutive frames; this threshold was empirically determined using the non-motile *ΔpilA* strain as a negative control. The motile population fraction was calculated by dividing the number of motile cells by the total number of tracked cells. Non-motile cells were excluded from scatter plot. Trajectory extraction, data analysis, and plotting were performed using custom Python scripts (version 3.9.16).

### Optical tweezer assay

Optical tweezer experiments were performed using a dual-trap optical tweezers system (C-Trap, LUMICKS, Amsterdam) controlled by Bluelake software. Purified T4P variants (WT, R30E, and G75L) were adjusted to a concentration of 0.1-0.2 mg/mL. Carboxylate-modified polystyrene beads (2 µm diameter; Invitrogen, #F8826) were diluted 1:2000 in PBS, and sonicated for 15 second. These beads were used to tether the purified pili via non-specific hydrophobic interactions. The microfluidic flow cell was operated in the laminar flow regime, preventing mixing between adjacent channels. The first channel was passivated with 1% casein according to the manufacturer’s protocol. Following passivation, the beads, pili, and PBS were introduced into the first, second, and third channels, respectively, under continuous low-velocity flow.

Two beads were optically trapped, and the flow was stopped to calibrate trap stiffness. Calibration was performed via power spectral density analysis of the thermal fluctuations of the trapped beads. Trap stiffness was adjusted to 0.68-0.71 pN/nm by tuning the overall power while maintaining the trapping laser at 100%, after which the force was zeroed.

The optically trapped beads were moved to the second channel to capture pili. Once pili bundles were tethered between the two beads, the assembly was gently transferred to the third channel (containing only PBS). The flow was stopped to perform force measurements. One optical trap was displaced slowly to stretch the pili. This progressive extension allowed individual filaments within the bundle to detach sequentially from the beads until complete rupture occurred.

During pulling, raw force and distance data were continuously acquired. For data analysis, the low-frequency force channel (“Force LF”) was extracted directly from the C-Trap .h5 files using the LUMICKS Pylake library^70^. This provided data downsampled to 78 Hz for robust mechanical analysis. Force-distance (f-d) curves were segmented into three distinct mechanical regimes. Phase I represents the initial extension period, characterized by a lack of measurable force response. As extension progresses, the system enters Phase II, exhibiting a non-linear force-extension relationship. The transition from this non-linear regime to Phase III (a regime of linear elastic response) was determined through targeted slope analysis. Specifically, an onset coordinate was manually identified and paired with a terminal coordinate recorded immediately prior to the abrupt force drop to 0 pN, which corresponds to a detachment event. To establish the linearity of Phase III, the effective spring constant (*k*) was determined by calculating the gradient (change in force / change in distance) between these bounding coordinates. This linear regime assignment was validated by ensuring that intermediate data points within the selected interval closely aligned with the calculated slope. Data extraction, analysis, and plotting were performed using custom Python scripts (version 3.9.16). A total of *n* = 20, 29, and 32 independent pili pulling events were analyzed for the WT, R30E, and G75L variants, respectively.

### Statistical testing

All statistical analyses were performed using Python. To assess differences across multiple independent groups, we used the non-parametric Kruskal-Wallis H-test. When the Kruskal-Wallis test indicated global statistical significance, Dunn’s post-hoc test was subsequently applied to identify specific pairwise differences between groups. To control the family-wise error rate, the resulting *P* values from the pairwise comparisons were adjusted using the Holm-Bonferroni correction method. Adjusted *P* values less than 0.05 were considered statistically significant. In the analyses, \*\**P* < 0.01, \*\*\**P* < 0.001.

