## Supplementary Data Figures and Tables for "A structural trade-off balances mechanical resilience and supramolecular adaptability in type IV pili"

### Supplementary Data 1

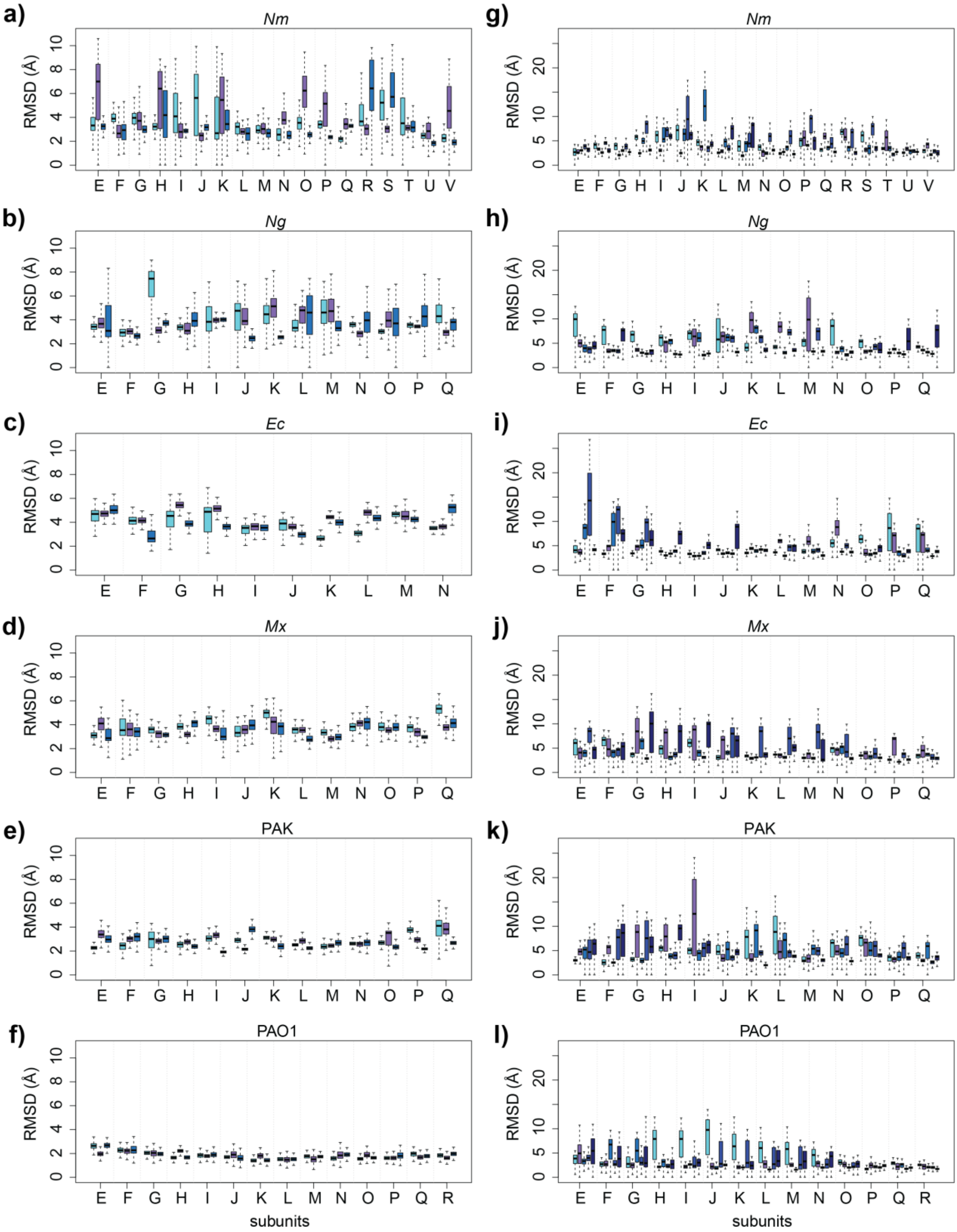


**Fig S1. Root mean square deviations (RMSD) of individual subunits across equilibrium and steered molecular dynamics (MD) simulations.**

RMSD values were calculated for the $C\alpha$ atoms of the pilin subunits relative to their equilibrated conformations for *N. meningitidis* (*Nm*; **a, g**), *N. gonorrhoeae* (*Ng*; **b, h**), *E. coli* (*Ec*; **c, i**), *M. xanthus* (*Mx*; **d, j**), and *P. aeruginosa* strains PAK (**e, k**) and PAO1 (**f, l**). Panels in the left column (**a-f**) display data from classical equilibrium MD simulations, whereas panels in the right column (**g-l**) display data from steered MD (SMD) simulations. Bar graphs represent the RMSD values for each subunit aggregated from three independent classical MD replicates and five independent SMD replicates.

#### Supplementary Data 2

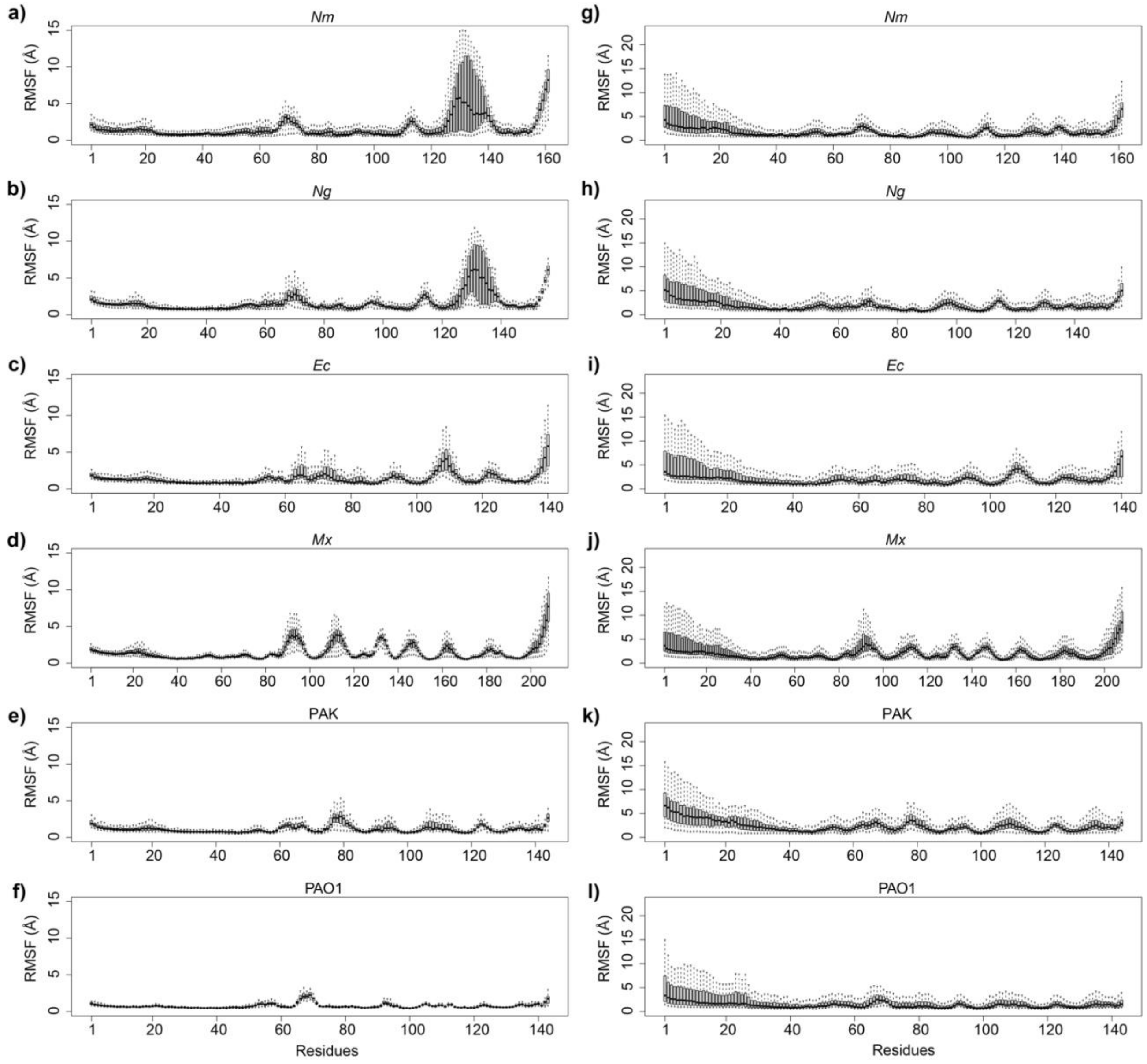


**Fig S2. Root mean square fluctuations (RMSF) of individual residues during classical and steered molecular dynamics (MD) simulations.**

RMSF values were calculated for the *Cα* atoms of each residue relative to the average simulation conformation. Data are shown for all bulk pilin subunits of *N. meningitidis* (*Nm*; **a, g**), *N. gonorrhoeae* (*Ng*; **b, h**), *E. coli* (*Ec*; **c, i**), *M. xanthus* (Mx; **d, j**), and *P. aeruginosa* strains PAK (**e, k**) and PAO1 (**f, l**). Panels in the left column (**a-f**) display data from classical equilibrium MD simulations, whereas panels in the right column (**g-l**) display data from steered MD (SMD) simulations. Box plots represent the RMSF values for a given residue across all bulk chains from three independent classical MD replicates and five independent SMD replicates.

### Supplementary Data Tables

#### Supplementary Data Table 1: Strains used in this study.

| **Name and relevant genotype** | **Source / Reference** | **Identifier** |
|---|---|---|
| XL10 Gold | Agilent | Na |
| *Escherichia coli* strain S17.1 (thi pro hsdR recA RP42(Tc::Mu)(Km::Tn7)) | Stratagene | Na |
| *Pseudomonas aeruginosa* PAO1 ΔfliC | (Bertrand et al, 2010)[1] | 177 |
| PAO1 Δ*fliC*Δ*pilT* | (Talà et al. 2019)[2] | 335 |
| PAO1 Δ*fliC*Δ*pilA* | This study | 1047 |
| PAO1 Δ*fliC*Δ*pilT*Δ*pilA* | This study | 1209 |
| PAO1 Δ*fliC*Δ*pilT*Δ*pilA*Δ*pilH* | This study | 1276 |
| PAO1 Δ*fliC pilA_E5L* | This study | 2453 |
| PAO1 Δ*fliC pilA_Y24T* | This study | 2454 |
| PAO1 Δ*fliC pilA_R30E* | This study | 2455 |
| PAO1 Δ*fliC pilA_R30A* | This study | 2634 |
| PAO1 Δ*fliC pilA_E49D* | This study | 2635 |
| PAO1 Δ*fliC pilA_E49R* | This study | 2636 |
| PAO1 Δ*fliC pilA_E49A* | This study | 2637 |
| PAO1 Δ*fliC pilA_I15A* | This study | 2785 |
| PAO1 Δ*fliC pilA_S35A* | This study | 2786 |
| PAO1 Δ*fliC pilA_Q23A* | This study | 2787 |
| PAO1 Δ*fliC pilA_E48A* | This study | 2789 |
| PAO1 Δ*fliC pilA_E48R* | This study | 2790 |
| PAO1 Δ*fliC pilA_G75A* | This study | 2791 |
| PAO1 Δ*fliC pilA_G75V* | This study | 2792 |
| PAO1 Δ*fliC pilA_G75L* | This study | 2793 |
| PAO1 Δ*fliC*Δ*pilT*Δ*pilH pilA_R30E* | This study | 2798 |
| PAO1 Δ*fliC*Δ*pilT*Δ*pilH pilA_G75L* | This study | 2860 |

**Supplementary Data Table 2: Plasmids used in this study.**

| Name and relevant genotype | Source / Reference | Identifier |
|---|---|---|
| pEX18Gm (Suicide vector based on pUC18, GmR, ColE1 ori (E. coli), oriT, sacB, lacZα) | (Hoang et al. 1998)[3] | Na |
| pEX100TAP (Suicide vector based on pUC19, AmpR, ColE1 ori (E. coli), oriT, sacB, lacZα) | (Schweizer & Hoang, 1995)[4] | Na |
| pEX18Gm_pilA_del | This study | pMK24 |
| pEX100TAP_pilH_del | (Barken et al.2008)[5] | 61 |
| pEX18Gm_pilA_E5L | This study | pCNT027 |
| pEX18Gm_pilA_Y24T | This study | pCNT028 |
| pEX18Gm_pilA_R30E | This study | pCNT029 |
| pEX18Gm_pilA_R30A | This study | pCNT040 |
| pEX18Gm_pilA_E49D | This study | pCNT041 |
| pEX18Gm_pilA_E49R | This study | pCNT042 |
| pEX18Gm_pilA_E49A | This study | pCNT043 |
| pEX18Gm_pilA_I15A | This study | pCNT051 |
| pEX18Gm_pilA_S35A | This study | pCNT052 |
| pEX18Gm_pilA_Q23A | This study | pCNT053 |
| pEX18Gm_pilA_E48A | This study | pCNT055 |
| pEX18Gm_pilA_E48R | This study | pCNT056 |
| pEX18Gm_pilA_G75A | This study | pCNT057 |
| pEX18Gm_pilA_G75V | This study | pCNT058 |
| pEX18Gm_pilA_G75L | This study | pCNT059 |

**Supplementary Data Table 3: Oligonucleotides used in this study.**

| Purpose | Name | Sequence |
|---|---|---|
| Generation of pMK24 | OL_PilA_KO_R | GTTATCACACATGAATCTCTCCGTTGATTATGTATAGG |
| | OL_PilA_KO_F | AGATTCATGTGTGATAACTAAGGTGATCGAAGG |
| | Xba1_PilA_F | CATGCCTGCAGGTCGACTCGGTAGCGCTTTCGAACAGC |
| | EcoR1_PilA_R | GCTATGACCATGATTACGCGGTGAAGCTCCTCGAC |
| Check pEX18 | seq_pEX18_F | CACTTTATGCTTCCGGCTCG |
| | seq_pEX18_R | TAAGTTGGGTAACGCCAGGG |
| Check pilA locus | seq_pilA_mutants_F | AGCGAGTCGAGCCGAACCAGAT |
| | seq_pilA_mutants_R | CGAGACCGTCGGTAGCGCTTTC |
| Common for all pilA mutants | OL_EcoRI_pilA_F | GCTATGACCATGATTACGTACCGCTACCCGCAGCCA |
| | OL_XbaI_pilA_R | CATGCCTGCAGGTCGACTGGTCGGAGATGCCTACAAAGAGCT |
| Generation of pCNT027 | OL_pilA_E5L_F | CGCAACCACGATCATCAGCAGGATCAAGGTAAAGCCTTTTTGAGCTTTCAT |
| | OL_OL_pilA_E5L_R | ATGAAAGCTCAAAAAGGCTTTACCTTGATCCTGCTGATGATCGTGGTTGCGATCATCG |
| Generation of pCNT028 | OL_pilA_Y24T_F | AACGCGCAACATAGTTCTGGGTCTGGGGAATGGCAATTGCCG |
| | OL_pilA_Y24T_R | CGGCAATTGCCATTCCCCAGACCCAGAACTATGTTGCGCGTT |
| Generation of pCNT029 | OL_pilA_R30E_F | AGCGCCGAAGCACCTTCCGACTCCGCAACATAGTTCTGATACTGGGGAATG |
| | OL_pilA_R30E_R | CATTCCCCAGTATCAGAACTATGTTGCGGAGTCGGAAGGTGCTTCGGCGCT |
| Generation of pCNT040 | OL_pilA_R30A_R | TATCAGAACTATGTTGCGGCCTCGGAAGGTGCTTCGGCG |
| | OL_pilA_R30A_F | AGCGCCGAAGCACCTTCCGAGGCCGCAACATAGTTCTGATACTGGGGAAT |
| Generation of pCNT041 | OL_pilA_E49D_R | CGCTGAAGACCACTGTTGAAGACTCGCTGTCGCGTGGAATTG |
| | OL_pilA_E49D_F | GCAATTCCACGCGACAGCGAGTCTTCAACAGTGGTCTTCAGCGGG |
| Generation of pCNT042 | OL_pilA_E49R_R | CGCTGAAGACCACTGTTGAACGCTCGCTGTCGCGTGGAATTG |
| | OL_pilA_E49R_F | GCAATTCCACGCGACAGCGAGCGTTCAACAGTGGTCTTCAGCGGG |
| | OL_pilA_E49A_R | CGCTGAAGACCACTGTTGAAGCCTCGCTGTCGCGTGGAATTG |

| | | |
|---|---|---|
| Generation of pCNT043 | OL_pilA_E49A_F | GCAATTCCACGCGACAGCGAGGCTTCAACAGTGGTCTTCAGCGGG |
| Generation of pCNT051 | OL_pilA_I15A_R | CGTGGTTGCGATCATCGGTGCCCTGGCGGCAATTGCCATTCC |
| | OL_pilA_I15A_F | GGAATGGCAATTGCCGCCAGGGCACCGATGATCGCAACCACGATC |
| Generation of pCNT052 | OL_pilA_S35A_R | TTGCGCGTTCGGAAGGTGCTGCCGCGCTGGCGACGATCAAC |
| | OL_pilA_S35A_F | CGGGTTGATCGTCGCCAGCGCGGCAGCACCTTCCGAACGCGC |
| Generation of pCNT053 | OL_pilA_Q23A_R | TGGCGGCAATTGCCATTCCCGCCTATCAGAACTATGTTGCGCGTTCGG |
| | OL_pilA_Q23A_F | CGCGCAACATAGTTCTGATAGGCGGGAATGGCAATTGCCGCC |
| Generation of pCNT055 | OL_pilA_E48A_R | AACCCGCTGAAGACCACTGTTGCCGAGTCGCTGTCGCGTGGAATT |
| | OL_pilA_E48A_F | ATTCCACGCGACAGCGACTCGGCAACAGTGGTCTTCAGCGGGTTG |
| Generation of pCNT056 | OL_pilA_E48R_R | AACCCGCTGAAGACCACTGTTCGCGAGTCGCTGTCGCGTGGAATT |
| | OL_pilA_E48R_F | ATTCCACGCGACAGCGACTCGCGAACAGTGGTCTTCAGCGGGTTG |
| Generation of pCNT057 | OL_pilA_G75A_R | GCTTCTACTGCGACCGAAACATATGTCGCCGTCGAGCCGGATGCCAACAAG |
| | OL_pilA_G75A_F | TTGTTGGCATCCGGCTCGACGGCGACATATGTTTCGGTCGCAGTAGAAGC |
| Generation of pCNT058 | OL_pilA_G75V_R | GCTTCTACTGCGACCGAAACATATGTCGTGGTCGAGCCGGATGCCAACAAG |
| | OL_pilA_G75V_F | TTGTTGGCATCCGGCTCGACCACGACATATGTTTCGGTCGCAGTAGAAGC |
| Generation of pCNT059 | OL_pilA_G75L_R | GCTTCTACTGCGACCGAAACATATGTCCTGGTCGAGCCGGATGCCAACAAG |
| | OL_pilA_G75L_F | TTGTTGGCATCCGGCTCGACCAGGACATATGTTTCGGTCGCAGTAGAAGC |
